# Remotely detected plant function in two midwestern prairie grassland experiments reveals belowground processes

**DOI:** 10.1101/2021.09.08.459443

**Authors:** Jeannine M. Cavender-Bares, Anna K. Schweiger, John A. Gamon, Hamed Gholizadeh, Kimberly Helzer, Cathleen Lapadat, Michael D. Madritch, Philip A. Townsend, Zhihui Wang, Sarah E. Hobbie

**Affiliations:** Department of Ecology, Evolution, and Behavior, University of Minnesota, Saint Paul, MN 55108; Institut de recherche en biologie végétale & Département de sciences biologiques, Université de Montréal, Montréal, QC, Canada; Department of Geography, Remote Sensing Laboratories, University of Zurich, Switzerland; School of Natural Resources, University of Nebraska Lincoln, Lincoln, NE; Departments of Earth & Atmospheric Sciences and Biological Sciences, University of Alberta, Edmonton, AB, Canada; Center for Applications of Remote Sensing, Department of Geography, Oklahoma State University, Stillwater, OK 74078; Department of Biology, Appalachian State University, Boone, NC; Department of Forest and Wildlife Ecology, University of Wisconsin-Madison, Madison, WI

**Keywords:** Biodiversity, enzyme activity, hyperspectral data, microbial biomass, plant traits, remote sensing, vegetation chemistry

## Abstract

Imaging spectroscopy provides the opportunity to incorporate leaf and canopy optical data into ecological studies, but the extent to which remote sensing of vegetation can enhance the study of belowground processes is not well understood. In grassland systems, aboveground and belowground vegetation quantity and quality are coupled, and both influence belowground microbial processes and nutrient cycling, providing a potential link between remote sensing and belowground processes. We hypothesized that ecosystem productivity, and the chemical, structural and phylogenetic-functional composition of plant communities would be detectable with remote sensing and could be used to characterize belowground plant and soil processes in two grassland biodiversity experiments—the BioDIV experiment at Cedar Creek Ecosystem Science Reserve in Minnesota and the Wood River Nature Conservancy experiment in Nebraska. Specifically, we tested whether aboveground vegetation chemistry and productivity, as detected from airborne sensors, predict soil properties microbial processes and community composition. Imaging spectroscopy data were used to map aboveground biomass and green vegetation cover, functional traits and phylogenetic-functional community composition of vegetation. We examined the relationships between the image-derived variables and soil carbon and nitrogen concentration, microbial community composition, biomass and extracellular enzyme activity, and soil processes, including net nitrogen mineralization. In the BioDIV experiment—which has low overall diversity and productivity despite high variation in each—belowground processes were driven mainly by variation in the amount of organic matter inputs to soils. As a consequence, soil respiration, microbial biomass and enzyme activity, and fungal and bacterial composition and diversity were significantly predicted by remotely sensed vegetation cover. In contrast, at Wood River, where plant diversity and productivity were consistently higher, remotely sensed functional, chemical and phylogenetic composition of vegetation predicted belowground extracellular enzyme activity, microbial biomass, and net nitrogen mineralization rates, while aboveground biomass did not. The strong, contrasting associations between the quantity and chemistry of aboveground inputs with belowground soil processes and properties provide a basis for using imaging spectroscopy to understand belowground processes across productivity gradients in grassland systems. However, a mechanistic understanding of how above and belowground components interact among different ecosystems remains critical to extending these results broadly.

## Introduction

Monitoring biodiversity and understanding its consequences for ecosystem functions and global processes are critical challenges in the face of rapid global change. Remote sensing has proven useful for observing ecosystem functions such as total biomass production (Gamon & Qiu, 1999; Williams et al., 2020) across spatial scales and temporal resolutions because reflectance and absorption of light by vegetation canopies is strongly influenced by their structural, biochemical, physiological, and phenological characteristics (Ustin & Gamon, 2010; Schmidtlein et al., 2012). The functional variation in vegetation can be detected using imaging spectroscopy (aka hyperspectral imagery) (Asner & Martin, 2009; Asner et al., 2011; Wang et al., 2016; Schneider et al., 2017; Schweiger et al, 2017; Wang et al., 2019; Williams et al. 2020), thus enabling hyperspectral remote sensing as an approach to test the drivers of ecosystem processes that can be detected aboveground.

In contrast to aboveground processes and attributes, detecting belowground processes remotely remains technically challenging and enigmatic. Nevertheless, the composition, function and diversity of plant assemblages are well known to affect belowground processes (Hooper & Vitousek, 1998; Eviner & Chapin, 2003; Meier & Bowman, 2008; Bardgett & van der Putten, 2014; Hobbie, 2015). Plants synthesize a wide variety of chemical and structural compounds to support physiological functions, and the abundance and chemical composition of plant tissues and plant exudates influence soil microbial diversity and abundance (Meier & Bowman, 2008; Hobbie, 2015). Given that plant reflectance spectra are aggregate indicators of chemistry, composition and abundance of plants within communities (Schmidtlein, 2005; Townsend et al., 2013; Serbin et al., 2014; Singh et al., 2015; Schweiger et al., 2018), it is appropriate to ask whether we can remotely sense attributes of vegetation that predict belowground processes (Madritch et al., 2014; Cavender-Bares et al., 2017).

Both quantity and quality of inputs from above to belowground may influence belowground processes. The total quantity of inputs to the soil from above and belowground primary productivity is expected to influence microbial processes by providing energy to fuel soil microbes (Zak et al. 1994; Hättenschwiler & Jørgensen, 2010). Thus, higher rates of plant inputs (above and belowground) to soils are often associated with higher rates of soil respiration, enzyme activity and nutrient mineralization (Zak et al., 1990; Cline et al., 2018).

The chemistry (“quality”) of organic matter inputs to the soil can also influence the abundance of different microbial organisms with contrasting metabolic capacities (Degens & Harris, 1997; Wardle et al., 2004; Cline et al., 2018). Variation in quality of inputs results from variation in plant foliar and root chemistry because live and dead plant parts include a diversity of organic molecules—such as soluble sugars, cellulose, hemicellulose, lignin, and tannins—which vary greatly in how readily they can be broken down by microbes (Degens & Harris, 1997; Meier & Bowman, 2008; Freschet et al 2012). Variation in plant nutrient concentrations thus influences organic matter decomposition and nutrient dynamics (Parton et al., 2007; Cornwell et al., 2008; Fornara et al., 2009).

Plant chemical diversity is in turn linked to physiological function associated with functional groups that often have a phylogenetic basis due to shared ancestry (Kothari et al., 2018; Cadotte et al. 2009), such as nitrogen fixation in legumes and C4 photosynthesis (Hooper & Vitousek, 1998; Craine et al., 2002). Plants can thus be meaningfully categorized into phylogenetic-functional groups (Kothari et al., 2018) that have a high potential to be remotely detected (Cavender-Bares et al., 2016; Schweiger et al., 2018; Wang et al., 2019; Meireles et al. 2020) as a basis to reveal belowground processes (Madritch et al., 2014). Even differentiation between monocots and dicots can be informative. For example, despite high variability, inputs from forbs and legumes tend to be characterized by high nitrogen content and soluble sugars (Adams et al., 2016), which may favor microorganisms with high hydrolytic capacity. In contrast, inputs from grasses (monocots), particularly C4 grasses, tend to be characterized by high cellulose and hemicellulose but low nitrogen content (Craine et al., 2001), which may favor microorganisms with high cellulose degrading activity. Thus, variation in plant composition—measured in terms of plant chemistry and function or in terms of variation in phylogenetic-functional group representation—is expected to influence microbial and soil processes, independent of the total quantity of inputs.

The capability of airborne imaging spectroscopy to accurately detect plant chemistry and productivity provides an important means to estimate both the quality and the quantity of aboveground inputs to the soil over large spatial scales (Kokaly et al., 2009; Asner et al., 2011; Serbin et al., 2014; Singh et al., 2015; Wang et al., 2019). Remotely sensed information may be useful in understanding soil processes both because aboveground inputs are significant sources of organic matter to soils and/or because above and belowground attributes (total net primary productivity [NPP], and chemical constituents) may be coupled. For example, in aspen (*Populus tremuloides*) stands, variation in canopy chemistry among genotypes—including condensed tannins, lignin, and nitrogen concentrations—was associated with foliar spectra and correlated with belowground processes (Madritch et al., 2014). In Hawaii, airborne spectroscopy detected the major source of variation in canopy nitrogen associated with both planted native nitrogen fixing trees (*Acacia koa*) and invading N fixers (*Myrica faya*) (Asner et al., 2008; Vitousek et al., 2009), which alter chemical inputs to soil microorganisms and influence soil processes (Vitousek, 2004). Unlike in these forest systems, organic matter inputs to soils in many grasslands are dominated by belowground productivity (Hui & Jackson 2006).

Nevertheless, aboveground processes such as photosynthesis are essential to providing belowground resources in grasslands, and the coupling of the drivers of aboveground primary production (quantity and chemistry) to those belowground could enable prediction of belowground processes and properties using remote sensing.

Here we investigated the extent to which we could remotely sense aboveground productivity, plant function, phylogenetic-functional group composition and spectral diversity at the scale of individual plant communities to predict belowground microbial (fungal and bacterial) biomass, composition, diversity and extracellular enzyme activity; net nitrogen mineralization rates; soil respiration; and soil carbon and nutrient concentrations. This work was conducted at the long-term prairie diversity experiment (BioDIV) at the Cedar Creek Ecosystem Science Reserve (CCESR) in central Minnesota (Tilman et al., 2001) and a more recent prairie diversity experiment at Wood River in central Nebraska established by the Nature Conservancy (TNC) (Nemec et al., 2013). The two experiments occur on different parts of the diversity-productivity gradient found in natural, degraded and restored prairie systems (Jelinski et al. 2011), with BioDIV experiment falling at the lower end of the gradient and Wood River at the higher end.

Previous work in the BioDIV experiment at CCESR showed strong linkages between aboveground plant composition and diversity to soil carbon and nutrients and microbial diversity and composition (Zak et al., 2003; Waldrop et al., 2006; Fornara, DA et al., 2009; Steinauer et al., 2015; Cline et al., 2018; Yang et al. 2019). In particular, higher fungal richness was associated with increasing aboveground plant biomass—and thus total plant-derived substrate inputs to soils—while fungal community composition and soil carbon and nitrogen cycling and pools were associated with plant functional group diversity, and the relative abundance of C4 grasses and legumes (Fornara, et al., 2009; Cline et al., 2018). Linkages between above and belowground processes have not yet been examined in the Wood River experiment, nor have linkages between remotely sensed variables and belowground processes in either experiment.

Our primary goals were 1) to decipher the mechanisms linking above and belowground processes in two contrasting grassland systems that differ in productivity, diversity, and soil type, and 2) to test the extent to which remotely sensed aboveground productivity, chemical characteristics, and phylogenetic-functional community composition can reveal belowground processes, microbial diversity and soil attributes. In pursuing these goals, we examined how well remotely sensed imaging spectroscopy could characterize aboveground functional and chemical composition, diversity and productivity and the strength and nature of the linkages among above- and belowground components and processes.

By comparing two experimental prairie systems that differed in the degree and range of plant diversity and productivity, but had similar functional group composition, we sought to understand whether relationships determined in one system applied to another system and what aspects of the linkages between above- and belowground processes must be understood to use remote sensing as a means to predict changes in belowground properties and processes. We hypothesized that the ecosystem productivity and the types of morpho-physiological and chemical properties of vegetation that vary among phylogenetic-functional groups would influence belowground productivity, composition and diversity of microbial communities and drive belowground ecosystem processes. We also hypothesized that the two systems would differ in the relative importance of the quantity (aboveground productivity) and the quality (chemical composition and phylogenetic-functional group composition) of inputs to the soil in driving belowground processes. In BioDIV—where productivity was relatively low—we expected effects of substrate quantity to overwhelm those of substrate quality, whereas in Wood River— where productivity was comparatively high—we expected the effects of substrate quality to be more apparent (Fig. 1). In both systems, we predicted that plant diversity would have a direct influence on productivity but would influence belowground processes primarily through its influence on productivity and the chemistry of plant inputs to soils. We tested these hypotheses using structural equation models (SEMs).

**Figure 1.**
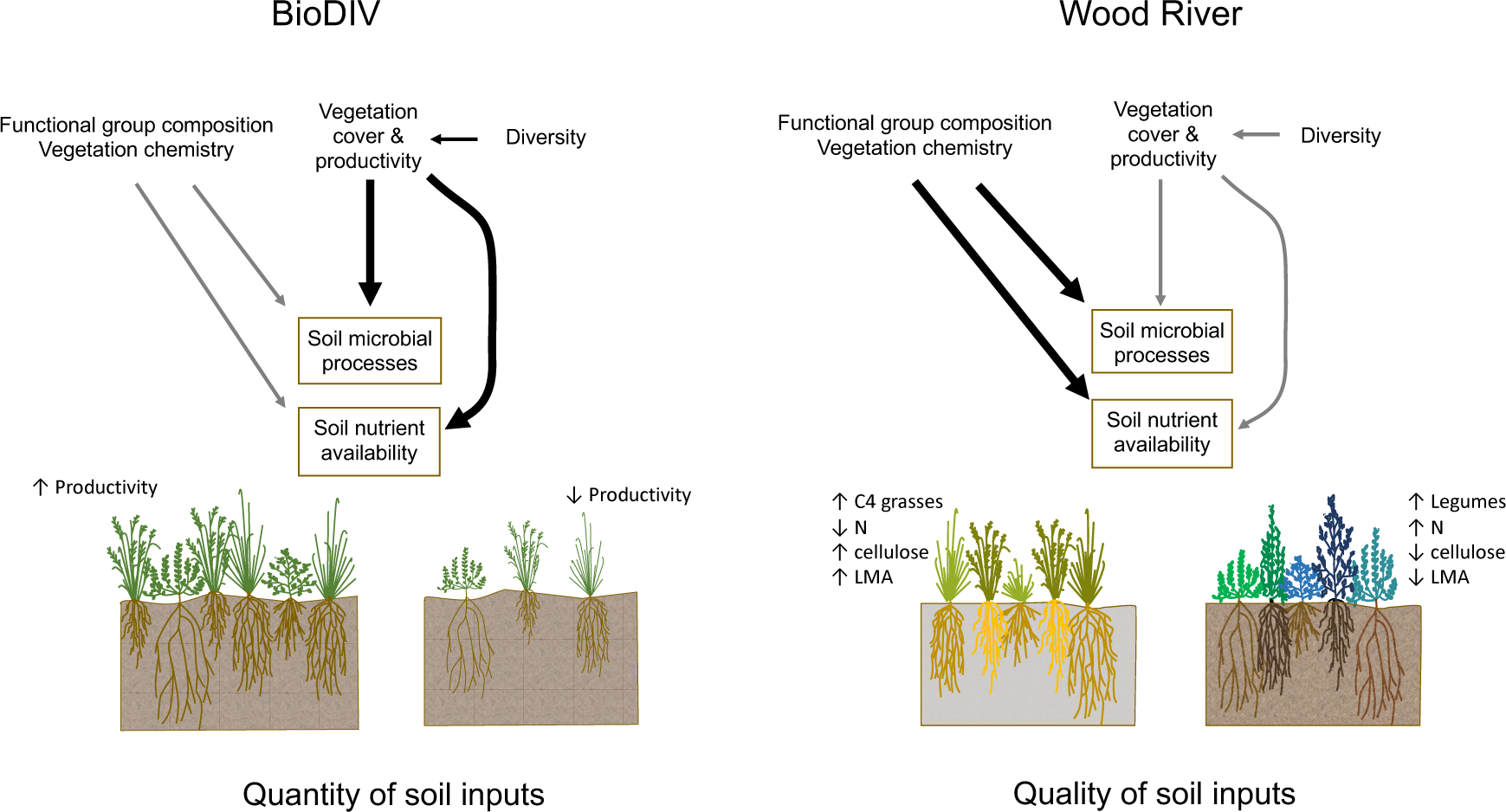
Both the quantity (productivity) and quality (phylogenetic, functional and chemical composition) of inputs from aboveground vegetation into the soil are important in influencing belowground microbial processes and nutrient cycling. However, the relative importance of these characteristics of inputs varies among systems. Shown are distinct hypotheses for how attributes of vegetation that can be remotely sensed influence and are thus useful in predicting soil microbial processes and nutrient availability in two contrasting prairie grassland diversity experiments. The BioDIV experiment at Cedar Creek varies in species density from one to 16 species per 81 m^2^, which has led to high variation among plots in annual productivity over time (from 25-283 g m^-2^). In this system, productivity (the quantity of inputs) is hypothesized to be the dominant driver of variation in belowground processes. The Wood River experiment has higher overall species density (one to 13 species per 1 m^2^ or 10-46 species per 1200 m^2^) and productivity (285-1079 g m^- 2^). As a consequence, the total quantity of inputs is less likely to be limiting, and functional group composition and the resulting variation in vegetation chemistry (quality of inputs) are hypothesized to play a more dominant role in driving belowground processes. These conceptual diagrams are the basis for our structural equation models presented in Figure 7.

Our study thus investigates how the quality and quantity of aboveground inputs influence belowground processes and attributes using imaging spectroscopic data to detect plant productivity, chemical and functional composition, and taxonomic, phylogenetic and functional diversity based on foliar chemical traits. Using this integrated approach, we determine the extent to which belowground processes can be inferred from imaging spectroscopy once mechanisms linking above and belowground processes have been characterized.

## Methods

### Study areas

The study was conducted at CCESR (East Bethel, Minnesota) in the long-term BioDIV experiment and at the Wood River prairie restoration experiment maintained by the Nature Conservancy (TNC) near Wood River, Nebraska. The sites differ in soil type, plot size, plot number, species composition, and the range of species of richness and aboveground productivity within plots.

The BioDIV experiment (N 45°24’11’’,W 93°11’21’’) was established in 1994 (Tilman, 1997; Tilman et al., 2001). Cedar Creek is located on a sandy glacial outwash plain and the soils where BioDIV is located are Typic Udipsamments (Grigal et al., 1974). Prior to the establishment of the experiment, topsoil was removed. In the full experiment, 1, 2, 4, 8, 16 or 32 perennial grassland species were planted from a pool of 34 species (eight species, each, of C4 grasses, C3 grasses, legumes, non-legume forbs; two species of woody plants) in a total of 342 9 x 9 m plots, separated by 1.5 meters of grass or bare soil. Of these, 154 have species composition and species richness levels maintained by annual weeding. The plots are also burned annually—such that annual aboveground biomass is a measure of aboveground productivity—with the consequence that litterfall has limited impact on soil organic matter buildup. We collected data in a subset of plots covering planted diversity levels ranging from 1 to 16 species per plot. The number of plots sampled and the specific measurements taken in each year are given in Table S1.

**Table 1.**
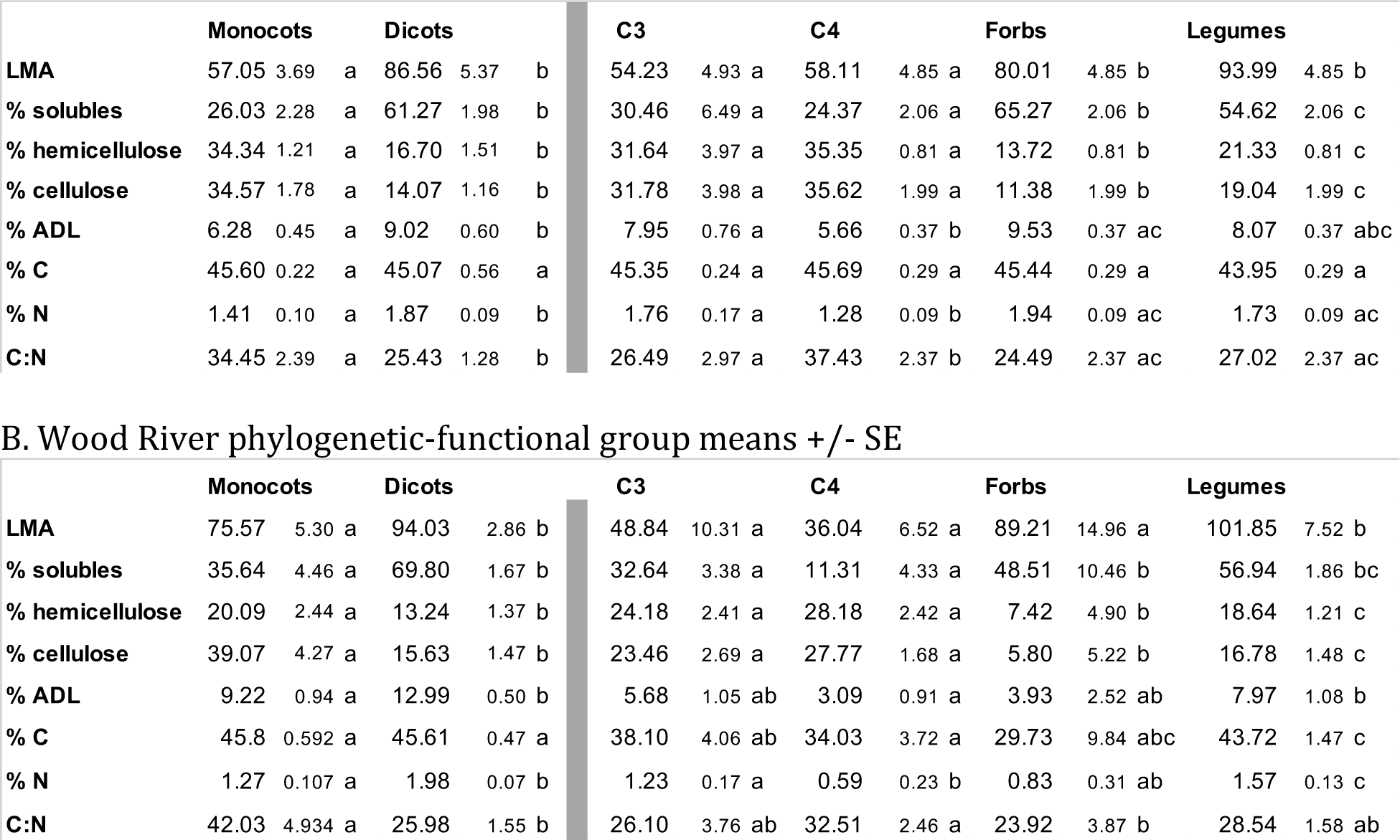
Phylogenetic-functional group means for chemical trait values derived from leaf level spectra for monocots, dicots, C3 grasses, C4 grasses, forbs and legumes. For BioDIV, trait values were averaged across all individuals of each of 29 species (Schweiger et al. 2018); for Wood River, trait values were averaged across all individuals of each of 49 species; Phylogenetic-functional group means were averages of species means. Significant differences between monocots and dicots (left) or among the C3 grass, C4 grass, forb and legume groups (right) based on Tukey’s HSD tests are shown with different letters, P cutoff=0.05.

The Wood River TNC experiment is a prairie restoration study along the Platte River, 10 km south of Wood River, Nebraska (N 40°44’37”, W 98°35’26”). The site is located within the Platte River floodplain and soils are of loamy alluvium or sandy alluvium parent material (NRCS, 2010). The site is characterized by a high water table, poor drainage, and high soil organic content (Jelinski & Currier, 1996) and was farmed as a corn and soybean rotation before being converted into an experimental prairie (Nemec et al., 2013). The TNC study area includes 36 60 × 60 m plots across two fields. The site thus has much larger plot sizes but fewer plots than BioDIV. As described in Nemec et al (2013) and Gholizadeh et al (2019), one set of 12 plots was seeded in 2010 (“young plots”) at low, medium and high species richness levels—with measured species richness per plot in 2017 ranging from 11 to 48 species. An earlier set of 24 plots was seeded in 2006 (“old plots”) with two diversity levels and two seeding rates. Plots are not weeded, and measured species richness in the old plots in 2017 varied from 28 to 46 species per plot. We collected leaf level spectral data, soil samples, and annual aboveground biomass in all of the young plots and in the 12 old plots with low seeding rate at both low and high diversity levels. Our study thus includes a total of 24 60 x 60 m plots. Given the large size of the plots, and heterogeneity within them, including gradients in hydrology towards the Platte River, we sampled eight 6-m^2^ subplots within each plot for a total of 192 subplots. Like the BioDIV, experiment, the site is managed with prescribed fires, although less frequently, and was last burned in late March 2015, two years prior to our study.

Several previous studies at each site used the airborne imagery, some of the soil data, and some of the leaf functional trait data (Table S1). This study goes beyond previous work by measuring soil processes at Wood River, adding new microbial data to BioDIV, quantifying leaf level functional traits using spectroscopy and mapping these traits as well as phylogenetic-functional groups at the landscape scale in Wood River, and by examining the associations between remotely detected spectroscopic information from aboveground vegetation and belowground processes. In combination, these studies represent a synthesis of our efforts to decipher the spectral signatures of plants and how they relate to ecosystem processes, with a focus on links between above- and belowground states and processes.

### Soil sampling

In the BioDIV experiment, soil was sampled to quantify microbial respiration from year-long incubations, net nitrogen mineralization, extracellular enzyme activity, and soil carbon and nitrogen concentrations. Soil was sampled to a depth of 10 cm at 1 m from the plot edge in four corners as well as the center (2 cm diameter, 5 cores/plot) and combined into a single sample per plot. In July 2014, 35 plots were sampled. Additional cores were taken in early August 2014 and washed over a 1mm sieve to collect roots. In July 2015, 125 plots (including the 35 plots from the 2014 sampling) were sampled. Soil carbon and nitrogen concentrations were measured in 154 plots using the same sampling approach in 2014, 2015, and 2016; means across years are presented here.

In the Wood River experiment, in 2017 soil was sampled to a depth of 10 cm in the center of each 6-m^2^ subplot located along transects approximately 20-22 m from the western and eastern borders of each plot and starting 12, 24, 36 and 48 m from the southern borders (192 subplots in total). From two aggregated soil cores per subplot, soil carbon and nitrogen, extracellular enzyme activity, potential net mineralization rate, and microbial biomass were measured.

Soils were composited by plot (BioDIV) or subplot (Wood River) and sieved (2 mm).

Subsamples were dried (65°C, 48 hours), milled, and total soil carbon and nitrogen measured by combustion using a Costech ECS4010 element analyzer (Valencia, California, USA). Subsamples of 2015 fresh soils from BioDIV were transported on ice and stored at - 80°C for molecular characterization of bacteria and fungi. Note that soil analyses from BioDIV (2015) were presented previously in Cline et al. (2018). We include those data as part of new analyses here.

In both experiments, soils were analyzed for microbial biomass carbon using a chloroform fumigation direct extraction procedure (Brookes et al., 1985). Within one to two weeks after collection and storage at 4°C, subsamples of sieved soil were extracted with 0.5 M K2SO4 (unfumigated) or extracted following fumigation in a chloroform atmosphere for 3 days (fumigated). Total dissolved carbon in extracts was determined on a TOC/TN analyzer (Shimadzu TOC-V, Shimadzu Corporation, Kyoto, Japan). Soil microbial biomass carbon and nitrogen were calculated as the difference between extractable carbon and nitrogen in the fumigated and unfumigated samples.

### Soil process rates

Total soil respiration rate was measured in the BioDIV experiment in 2015 as the accumulation of CO2 from 50 g of soil in airtight 1 L Mason jars during 24-48 h intervals on 16 dates throughout a year-long aerobic incubation (Cline et al., 2018). Cumulative carbon respired (mg CO2-C [g soil]^-1^) was calculated as the sum of the average respiration rate between adjacent measurement dates multiplied by the time interval between measurements and divided by the initial soil mass. We calculated net nitrogen mineralization rates (μg N [g soil]^-1^ d^-1^) as the difference between initial and final 2 M KCl-extractable concentrations of nitrogen as NH_#_ and NO_&_ (Horwáth, 2003) from 30-d incubations of field-moist soils in the laboratory, measured using microplate salicylate and sulfanilamide methods, respectively, on a BioTek Synergy H1 microplate reader (Winooski, Vermont, USA).

### Root chemistry

Roots were oven-dried at 65 °C for biomass determination. Root carbon fractions (cell solubles, hemicellulose and bound proteins [hemicellulose, hereafter], cellulose, and acid nonhydrolyzable residue, a measure of lignin plus recalcitrants [lignin, hereafter]) were determined with sequential digestion using an ANKOM fiber analyzer (ANKOM Technology, Macedon, New York, USA). Carbon and nitrogen concentration (% dry mass) were determined using combustion–reduction elemental analysis (TruSpec CN Analyzer; LECO, St. Joseph, Michigan, USA).

### Fungal and bacterial sampling

Fungal and bacterial composition and diversity were only determined in the BioDIV experiment. Soil samples were collected for the same 35 plots used for other analyses in 2014 and DNA extracted using the Fas-tDNA SPIN Kit (MP Biomedical, Solon, Ohio, USA). Fungal DNA analyses were reported in Cline et al (2018). Briefly, polymerase chain reaction (PCR) amplification of the ITS1 gene region was conducted using primers ITS1F and ITS2 (Smith and Peay 2014). Sequencing was performed on the MiSeq platform (Illumina, San Diego, California, USA) with 250 paired-end reads at West Virginia University’s Genomic Core Facility. For bacterial DNA analyses, which were not previously reported, 16S rRNA genes were amplified by PCR and sequenced at the West Virginia Genomics Core Facility in Morgantown, West Virginia, for sequence analysis with the Illumina MiSeq platform (Illumina, San Diego, California, USA) with 250 paired-end reads at West Virginia University’s Genomic Core Facility. Detailed methods are provided in the supplemental material (Appendix A). Aligned and screened sequences were clustered at 97% similarity and operational taxonomic unit (OTU) classification of unique sequences was completed using the Ribosomal Database Project (RDP) taxonomic database (release 9; Cole et al. 2009) and rarified to account for variation in the number of sequence reads per sample. Fastq files are stored with NCBI (SRA accession: 108802). Total OTU richness and inverse Simpson diversity are reported for fungi and bacteria; fungal OTU richness was previous published (Cline et al 2018). Non-metric multidimensional scaling (NMDS) ordination analyses were calculated for the fungal communities and bacterial communities within each plot using *metaMDS* in the vegan package in R (Oksanen et al., 2018) to determine the microbial community structure of the BioDIV plots.

### Extracellular microbial enzyme activity

In both the BioDIV and Wood River sites, we estimated the hydrolytic and oxidative (lignolytic) enzyme activity of soil communities using extracellular enzyme assays as described in Cline et al. (2018). We measured activity of *α*-glucosidase (AG, EC 3.2.1.20), *β*- 1,4-glucosidase (BG, EC 3.2.1.21), cellobiohydrolase (CBH, EC 3.2.1.91), *β*-1,4-xylosidase (BX, EC 3.2.1.37), and N-acetyl-*β*-glucosaminidase (NAG, EC 3.1.6.1), using methylumbellyferyl (MUB)-linked substrates (German et al., 2011). A 25-mM L-dihydroxy-phenylalanine substrate was used to assay phenol oxidase (PO, EC 1.10.3.2) and peroxidase (PX, EC 1.11.1.7) activity. Hydrolytic enzyme potential was calculated as the sum of the activities of CBH, AG, BG, BX, and NAG, and oxidative activity was calculated as the sum of PX and PO.

### Aboveground biomass and species sampling

At the BioDIV site, we quantified plot productivity and the relative dominance of plant functional groups by collecting aboveground plant biomass (g m^-2^) within the plots that are maintained for species richness levels 1 to 16 (2014, n=121; 2015, n=154; 2016, n=154) in a 9 m x 6 cm strip clip in late-July of each year. These plots include those in which soils were sampled. Aboveground biomass was sorted by plant species, dried at 60°C for 48 h and weighed. Aboveground biomass was assigned to 79 plant species and four plant phylogenetic-functional groups, including C3 grasses, C4 grasses, forbs and legumes, as well as the two major phylogenetic groups, monocots and dicots. We quantified total aboveground biomass of each plot, as well as the relative proportion of biomass (i.e., relative dominance) made up by individual plant functional groups. Vegetation cover was estimated using the point-quadrat method in 35 plots in 2014 (the same plots in which soil samples were collected that year).

At the Wood River site, we sampled aboveground biomass and percent cover of species within two 0.5 m^2^ areas within each subplot. These metrics were used to determine species richness and abundance at the subplot scale. In addition, we recorded presence of species along two 1-m wide transects, each centered approximately 20 m from the eastern and western borders of the plots. These data were used to determine species richness at the whole-plot scale.

### Foliar sampling

At the BioDIV site, spectral sampling at the leaf level, the collection of foliar samples, chemical assays and the prediction of foliar traits from spectra were described in detail in Schweiger et al. (2018). An SVC full-range (400-2500 nm) portable spectrometer (Spectra Vista Corporation, Poughkeepsie, New York, USA) with a leaf clip and tungsten-halogen light source (LC-RP PRO; Spectra Vista Corporation) was used to obtain foliar spectra and develop leaf trait models. Leaf-tissue samples of 130 individuals from 62 species were collected together with leaf spectra in the summers of 2015 and 2016 at the CCESR, flash frozen in liquid nitrogen and stored at -80 °C for subsequent pigment analysis. An additional set of 130 samples of 61 species were spectrally sampled and oven-dried at 65°C for chemical analyses of non-labile constituents. Pigment concentration and area-based pigment content was determined on flash-frozen samples using high-performance liquid chromatography (HPLC, Agilent 1200 Series; Agilent Technologies, Santa Clara, California, USA). The pigments included chlorophyll *a*, chlorophyll *b*, one carotene pigment (β- carotene), and five xanthophylls (lutein, zeaxanthin, violaxanthin, antheraxanthin, and neoxanthin). The total carotenoid pigment pool was calculated as the sum of five xanthophyll pigments plus β-carotene (Croft and Chen, 2018). Carbon fraction concentrations (% dry mass) were determined with sequential digestion, as described for roots above, for soluble cell contents, hemicellulose, cellulose, and lignin. Carbon and nitrogen concentration (% dry mass) were determined using combustion–reduction elemental analysis (TruSpec CN Analyzer; LECO, St. Joseph, Michigan, USA).

At the BioDIV site, leaf-level spectra were measured in four to eight 1-m^2^ subplots per plot, as described in Schweiger et al. (2018). At the Wood River site, leaf-level spectra were collected at regular intervals of 2 m along two transects starting at 2 m from the southern border and ending at 58 m, yielding a total of 29 individual plants measured per plot. Foliar trait models for all pigments, all carbon fractions, C and N were developed using partial least square regression (PLSR, Martens et al. 1983; Wold et al. 1983) implemented in the R package plsRglm (Bertrand et al., 2014) and are reported in Schweiger et al. (2018). Model performance ranged between 0.53 and 0.84 R^2^. The models were applied to all 1129 leaf-level spectra collected in the BioDIV experiment and all 1399 leaf-level spectra collected at Wood River. A PLSR model built from 892 grass and forb samples with concurrent SVC spectra collected during peak growing season in grassland ecosystems near Madison, Wisconsin was used to predict leaf mass per area (LMA, g m^-2^) for each individual plant sampled in the BioDIV (Wang et al 2019) and Wood River experiments.

Species mean leaf-level traits were scaled to the plot level based on relative biomass at the BioDIV experiment (Wang et al 2019). At Wood River, species mean traits were scaled to the subplot level using percent cover. The same was done for phylogenetic- functional groups. The whole plot- and whole subplot-level trait estimates, respectively, were matched to spectra extracted from airborne imagery to develop trait maps, as described below.

### Airborne spectroscopic data collection, trait and phylogenetic-functional group mapping

Imaging spectroscopy data from the Airborne Visible/Infrared Imaging Spectrometer -Next Generation (AVIRIS-NG, Hamlin et al., 2010) were collected at BioDIV with a spatial resolution of 0.9 m on the ground by the National Aeronautics and Space Administration (NASA) on August 25, 2014, August 30, 2015 and August 31, 2016 (Wang et al. 2019). The AVIRIS-NG reflectance data comprise 432 spectral bands and span 380 nm to 2510 nm with a spectral resolution of 5 nm. At Wood River, airborne imaging spectroscopy data were collected on August 23, 2017, with a spatial resolution of 1 m on the ground by an AISA Kestrel (Specim, Oulu, Finland) sensor operated by the University of Nebraska’s Center for Advanced Land Management Information Technologies (CALMIT) (Gholizadeh et al. 2019). The AISA Kestrel reflectance data comprise 178 spectral bands and span 400 nm to 1000 nm with a spectral resolution of approximately 3.5 nm (Golizadeh et al. 2020). We used partial least squares regression (PLSR, Wold et al., 1983) as implemented in the R package pls (Mevik et al. 2018) to model and predict biomass, functional group composition and foliar functional traits from imaging spectroscopy data, an approach widely used for the retrieval of plant traits and vegetation parameters (Asner & Martin, 2008; Serbin et al. 2014; Asner et al., 2015; Singh et al., 2015; Wang et al., 2019).

In BioDIV, plot-level trait data were linked to the spectroscopic images as reported in Wang et al (2019). In short, models for predicting functional traits from remotely sensed spectra were developed based on AVIRIS NG spectra and leaf trait models (Schweiger et al. 2018) scaled to the plot level with PLSR using plsregress in Matlab (The MathWorks, Inc., Natick, Massachusetts, USA) (Wang et al 2019). We modeled vegetation cover as a function of the mean spectral angles between soil and vegetation spectra per plot using logistic regression (see Serbin et al 2015) and predicted vegetation cover for all the pixels in the three images based on the resulting regression equation. We used a 7 × 7 pixel window centered in the 9 m × 9 m plots, and used mean spectra and trait estimates by plot as model inputs. Pixels with vegetation cover of less than 20% were excluded from analysis. For trait mapping, we applied PLSR coefficients to all pixels in the AVRIS NG images. For analysis with soils data, we used the predicted mean vegetation traits per plot for the central 7 × 7 pixel window and again excluded pixels with <20% cover.

In Wood River, similar to the approach used for modeling functional traits in BioDIV, we used plot- and subplot-level trait estimates and airborne spectra extracted from the same plots and subplots to model biomass and functional group composition and functional traits, including foliar soluble cell contents and the contents of carbon, nitrogen, hemicellulose and lignin with PLSR; here we used the R package pls (Mevik et al. 2018). For modeling biomass, phylogenetic-functional group composition and chemical traits, 25% of the full dataset was set aside for independent validation of the models; 75% was used to develop the models. Of this portion of the data, we used, depending on the number of observations, 75% or 80% of the data for model calibration and the rest for validation over 500 model runs. To avoid overfitting, the number of components used in PLSR was selected by minimizing the mean Root Mean Square Error (RMSE) and the cross-validated Prediction Residual Error Sum of Squares (PRESS) (Chen et al., 2004). The mean model coefficients with significant fits (P *≤* 0.05) were then applied to the spectral images to predict pixelwise biomass, functional group composition, and chemical traits. Model performance was assessed based on the R^2^ for linear regressions between measured and predicted values, the RMSE, % RMSE and model bias (see Table S2). Since the plots at the Wood River site had no significant soil exposure, pixels were not filtered based on vegetation cover.

### Diversity metrics

Phylogenetic species richness (PSR) (Helmus, 2007) was calculated using the phylogeny from Smith and Brown (Smith & Brown, 2018), pruned to include the measured species in the BioDIV experiment and the WoodRiver experiment separately. Absent species were added within the correct genus in a randomly assigned location using the congeneric merge function in pez in R (Pearse et al., 2015). Functional diversity was calculated using Scheiner’s functional dispersion [FD(qDTM)] (Scheiner et al. 2017) using biomass or percent cover (at Wood River) of each species within each plot to weight the species following Schweiger et al (2018). Remotely sensed spectral diversity using AISA airborne data in Wood River was calculated as Euclidean distances among vector-normalized reflectance values for all wavelengths among all pixels per plot, using the dist function in R.

### Structural equation modeling

The causal diagrams (Fig. 1) outline our hypotheses regarding how aboveground vegetation and soil processes are connected. We tested the degree of support for these hypotheses for the BioDIV and Wood River experiments, respectively, with SEMs as implemented in the R package lavaan (Rosseel, 2012). Measured variables were used as indicators of each of the constructs in the model. We used aboveground biomass as an indicator of vegetation quantity, and vegetation traits—including foliar chemical composition and specific leaf area (SLA)—and functional group abundance (proportion or biomass) as indicators of vegetation quality. Hydrolytic enzyme activity, microbial biomass and cumulative soil respiration were used as indicators of soil microbial activity. Net nitrogen mineralization rates and soil carbon and nitrogen concentration were used as indicators of soil nutrient availability. To avoid model overfitting, we selected those chemical vegetation traits and functional groups we considered to be the most important factors influencing vegetation quality in our study systems. Although more complex relationships are theoretically possible, we used linear specifications for our variables, which were also supported by the univariate regression analyses.

## Results

### Consistency between phylogenetic relationships, functional groups, and functional traits

In both BioDIV and Wood River, the functional groups that the diverse set of species represents largely correspond to monophyletic phylogenetic lineages, nested within the monocots or the dicots (eudicots)—the two major angiosperm lineages found in these ecosystems (Fig. 2). Within the monocots, C3 and C4 grasses in these experimental systems each fall within individual monophyletic lineages. The legumes and the forbs also each fall within distinct phylogenetic clades and—together with the small number of woody plants occasionally in the plots—are collectively nested within the dicot lineage. As a consequence, we refer to the functional groups as phylogenetic-functional groups.

**Figure 2.**
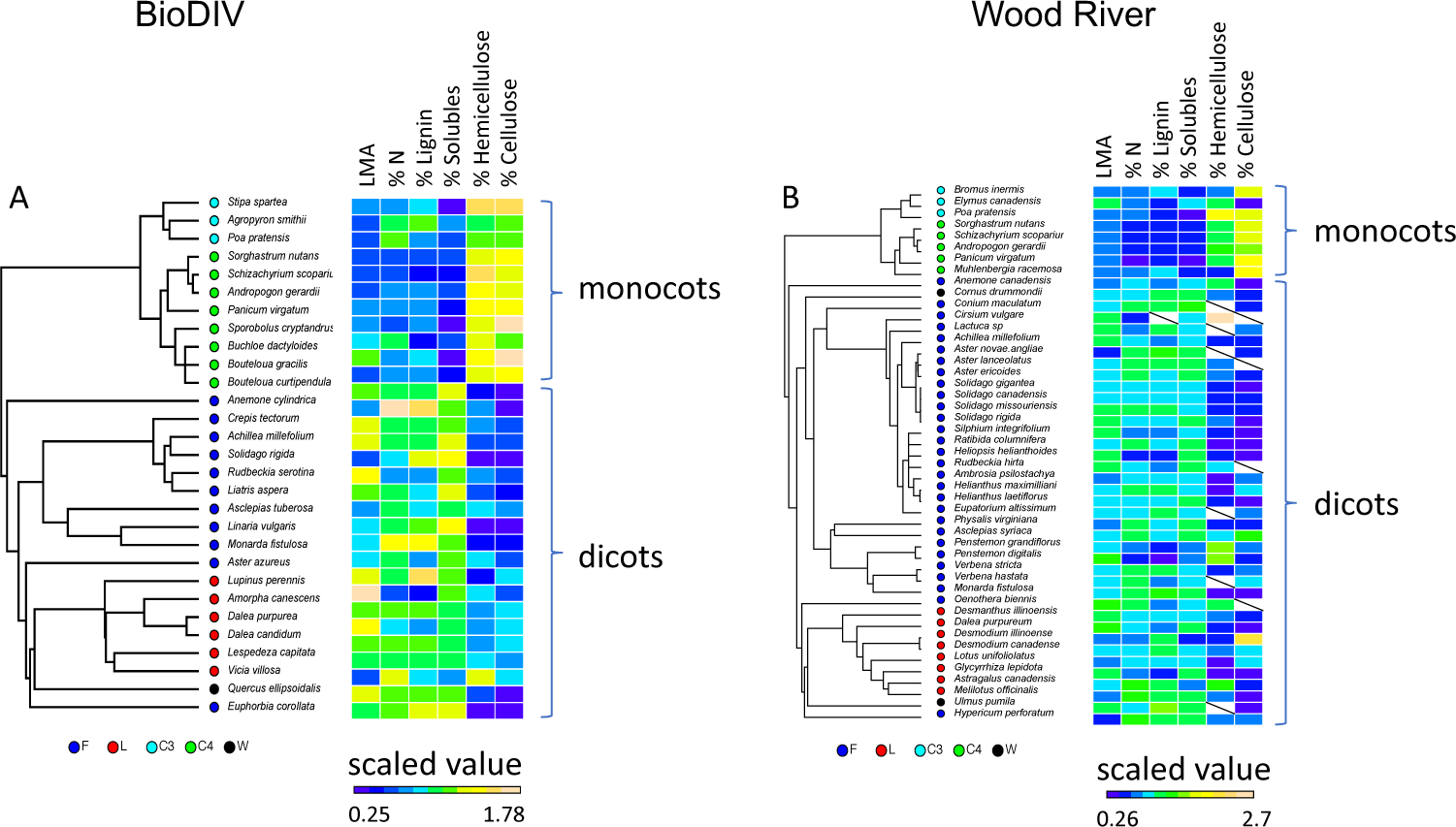
Phylogenetic relationships among species, their functional group categories (forbs, F; legumes, L; C3 grasses, C3; C4 grasses, C4; and woody species, W) and their foliar functional and chemical composition (leaf mass per area, LMA; % N, % lignin; % cell solubles; % hemicellulose, and % cellulose) in A) the BioDIV experiment at Cedar Creek and B) the Wood River experiment in Nebraska. Note that F, L and W groups are nested within the dicot lineage and C3 and C4 grasses are nested within the monocots. Functional trait values are scaled—centered to their means and divided by their standard deviations—and color-coded with blue colors indicating low values and yellow or salmon colors indicating high values. Missing values are indicated by a slash.

The morphological and chemical traits of species also tended to correspond to their phylogenetic-functional group identities. Differences in vegetation chemistry were consistently pronounced between monocots and dicots. In both systems, monocots (C3 and C4 grasses) had lower LMA, foliar nitrogen and lignin concentrations, lower cell soluble concentrations, and higher hemicellulose and cellulose concentrations compared to the forb and legume groups within the dicots (Table 1). The C4 grasses, in particular, tended to have high cellulose and hemicellulose concentrations. In contrast, the dicots tended to have lower foliar hemicellulose and cellulose concentrations and somewhat higher foliar nitrogen and lignin concentrations than the monocots (Fig. 2, and Table 1). Differences between groups within these two major lineages depended on the compound and to some extent, the site. Legumes had higher foliar nitrogen and lignin concentrations than the other groups at the Wood River site but only significantly higher foliar nitrogen and lignin than C4 grasses in BioDIV (Table 1). C4 grasses had lower concentrations of foliar nitrogen, lignin and solubles but higher concentrations of cellulose and hemicellulose than other functional groups, particularly at the Wood River site.

### Measured biodiversity-productivity relationships

In the BioDIV experiment, measured species richness (using 2014 data and counting only species that were originally planted in the experiment since the other ones are annually removed), strongly predicted aboveground biomass (R^2^=0.45, N=153, P<0.0001, Fig. 3A), as has been reported numerous times previously (Reich et al. 2012). Productivity was also significantly predicted by phylogenetic species richness (PSR), a measure of phylogenetic diversity that accounts for shared ancestry and evolutionary distance among species (R^2^=0.15, N=152, P<0.0001, Fig. 3B), and leaf level functional diversity [FD (qDTM)] calculated using mean traits per species and their abundances, based on biomass (R^2^=0.39, N=153, P<0001, Fig. 3C). The strengths of the relationships in different years were very similar. In the Wood River experiment, measured species richness was a weaker predictor of biomass than in the BioDIV experiment. At the subplot scale (6 m^2^), species richness explained 9% of the variation in aboveground biomass (R^2^=0.09, N=191, P<0.0001, Fig. 3D). At the plot scale, species richness explained 16% in aboveground biomass (R^2^=0.16, N=24, P=0.05, Fig. 3G). Phylogenetic species richness significantly but weakly predicted aboveground biomass at both the subplot scale (R^2^=0.07, N=191, P<0.0001, Fig. 3E) and the plot scale (R^2^=0.20, N=24, P=0.026, Fig. 3H). Abundance-weighted functional diversity [FD (qDTM)] also weakly predicted biomass at the subplot scale (R^2^=0.09, df=190, P<0.0001, Fig. 3F) and plot scale (R^2^=0.15, N=24, P=0.066, Fig. 3I).

**Figure 3.**
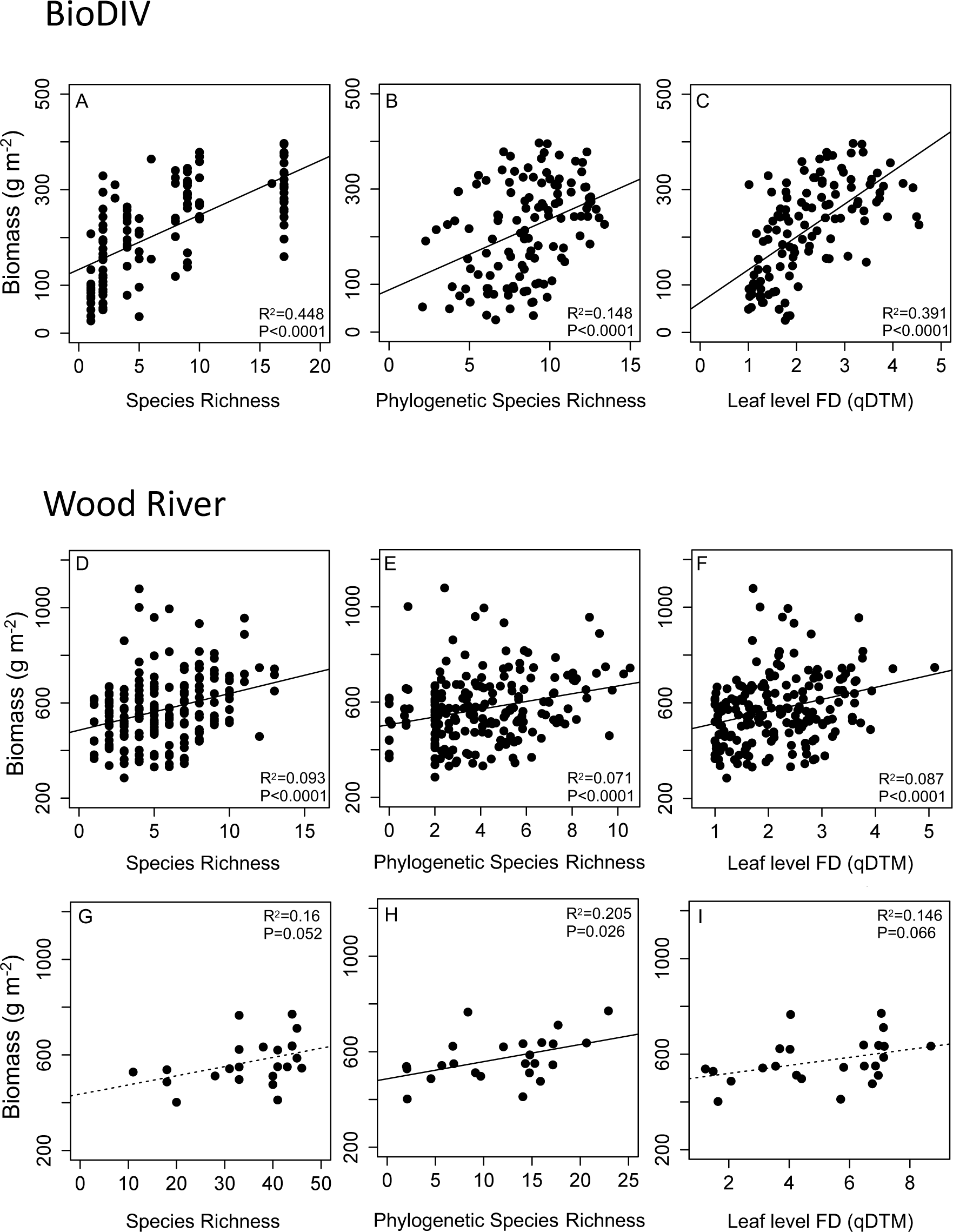
Biodiversity-Ecosystem Function relationships in the BioDIV and Wood River experiments. Shown are regressions between plant diversity metrics and annual aboveground biomass for BioDIV from 2014, N=124 (A-C) and Wood River at the subplot scale from 2017, N=192 (D-F) and plot scale, N=24 (G-I). Diversity metrics include measured species richness (SR), phylogenetic species richness (PSR) and abundance weighted functional diversity (FD (qDTM)) measured at the leaf level.

### Remote sensing of biomass (annual productivity)

Annual aboveground plant biomass, which is a measure of annual aboveground productivity in grasslands, varied considerably across the plots in the BioDIV experiment, ranging from approximately 19 to 740 g m^-2^. PLSR models developed for each of the three years using AVIRIS NG spectra predicted plot-level biomass well (Fig. 4A for 2014, Table S2). Independent validation results in 2014 (R^2^=0.75, 22 components, %RMSE= 12.05, P<00001), 2015 (R^2^=0.47, 18 components, %RMSE=16.4, P<00001) and 2016 (R^2^=0.64, 18 components, %RMSE=15.13, P<00001) were consistent (Table S2). Remotely sensed vegetation cover reported in Wang et al (2019) (R^2^=0.68, %RMSE=11)—based on the soil spectral angle calculated between the averaged soil spectrum and each pixel in the airborne image for the plot (Serbin et al. (2015)—also predicted biomass for the three years using linear regression (Fig. 4B, R^2^=0.63 for 2014, 0.70 for 2015, 0.55 for 2016, N=153, P<0.0001 all years, Table S2), serving as an accurate proxy of productivity in this system.

**Figure 4.**
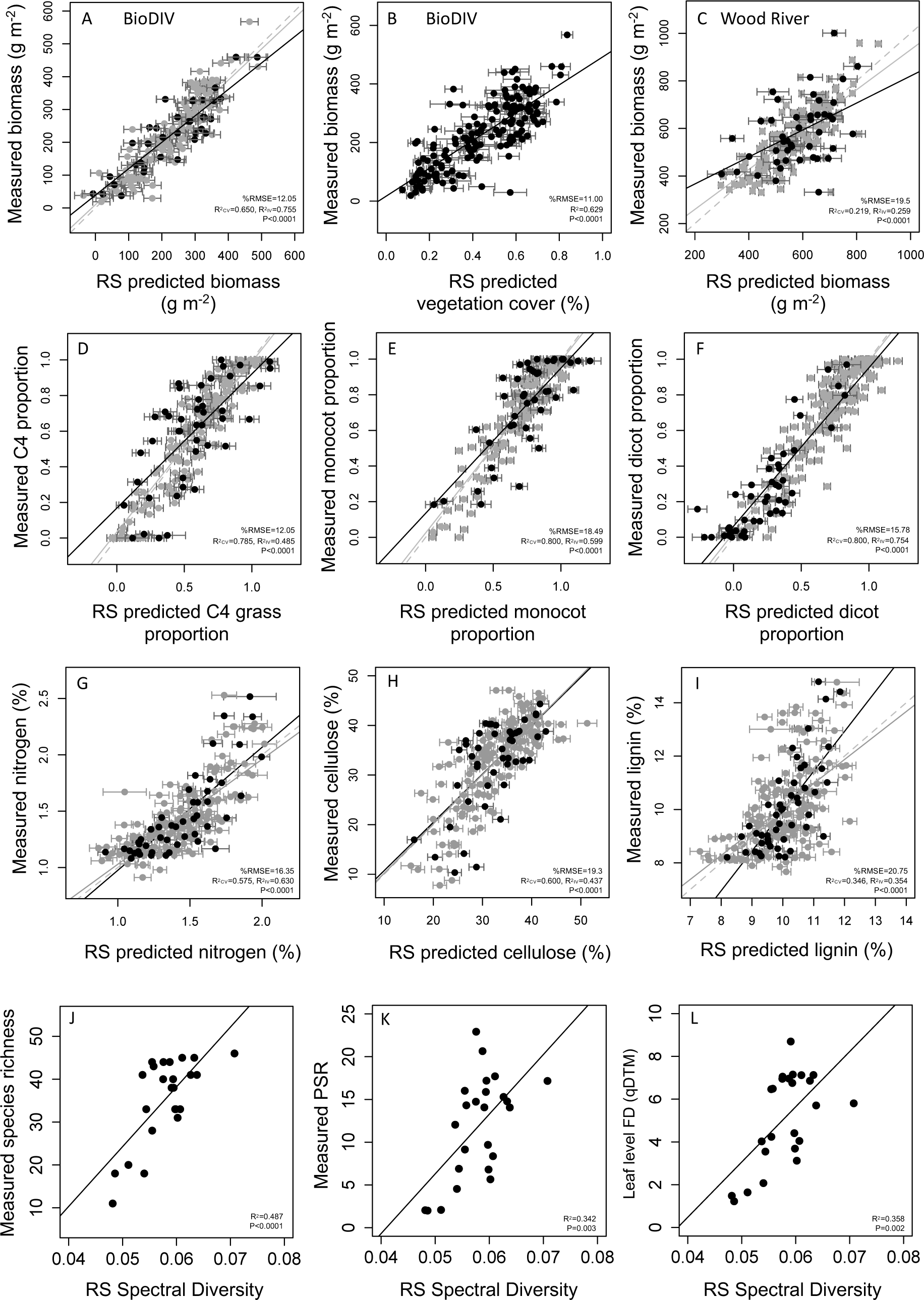
In the BioDIV experiment, biomass was accurately predicted by airborne spectroscopic imagery (AVIRIS NG) using PLSR (A) or remotely sensed vegetation cover (B). In the Wood River experiment, biomass was also predicted from spectroscopic imagery (AISA Kestrel) using PLSR at the subplot scale (C). Airborne data also predicted phylogenetic-functional group composition in the Wood River experiment—shown here for monocot (D), dicot (E) and C4 (F) proportions—and functional traits, including foliar nitrogen (G), cellulose (H) and lignin (I) concentrations. Both cross-validation (CV, gray circles) and independent validation (IV, black circles) %RMSE and R^2^ are shown for PLSR models. Note that the vegetation cover model was reported in Wang et al. 2019, with the %RMSE shown. These model predictions were applied to all BioDIV plots to predict measured biomass (B). In J-L, remotely sensed (RS) spectral diversity—the mean of the vector-normalized spectral distances among pixels within a plot—predicts measured species richness, phylogenetic species richness and abundance weighted functional diversity (FD(qDTM)) at the plot scale.

In the Wood River experiment, PLSR models also significantly predicted biomass at the subplot scale (R^2^=0. 26, 11 components, %RMSE=19.5, P<0.0001) (Fig. 4C, Table S2). When predicted values of biomass were applied to every pixel in the plots and averaged, predicted biomass at the plot scale was strongly associated with the average measured biomass in each plot (R^2^=0.76, N=24, P<0.0001, Fig. S3). In this system, vegetation cover was 100% throughout the experimental landscape (Gholizadeh et al., 2019) and remotely sensed vegetation cover did not predict biomass at either the subplot or plot scale.

### Detecting plant phylogenetic-functional groups from airborne spectroscopic imagery

In the Wood River experiment, the proportion of each phylogenetic-functional group within a subplot was significantly predicted from airborne imagery (Fig. 4D-F, Table S2), although with higher accuracy at broader phylogenetic levels—monocots, dicots— than for the groups within those. Independent validation results from PLSR models at Wood River using AISA Kestrel for monocot proportion (R^2^=0.55, 12components, %RMSE=18.49, P<0.0001), dicot proportion (R2=0.71, 11 components, %RMSE=15.78, P<0.0001), legume proportion (R^2^=0.24, 10 components, %RMSE=22.59, P<0.0001), forb proportion (R^2^=0.24, 11 components, %RMSE=19.12, P<0.001), C4 grass proportion (R^2^=0.43, 12 components, %RMSE=23.26, P<0.0001) and C3 grass proportion (R^2^ of 0.18, 11 components, %RMSE=22.48, P=0.004) showed that the proportion of these groups in each subplot could generally be predicted. In BioDIV, PLSR models predicting the biomass of individual phylogenetic-functional groups were inconsistent but most accurate and consistent for monocot biomass across years (Table S2). They were generally not able to predict the proportion of these groups likely due to the high exposed soil/vegetation cover in this experiment (Wang et al. 2018) and particularly to the limited number of plots with a high proportion of any given functional group that also had low exposed soil area.

### Functional trait models for Wood River

We developed PLSR models for functional traits to map functional traits using AISA data at Wood River (Table S2), following a similar approach as reported in Wang et al. (2019) for mapping traits in the BioDIV experiment from AVIRIS NG data but using %cover instead of biomass for calculating community weighted mean traits. Independent validation results for predicted foliar concentrations of nitrogen (R^2^=0.58, components=5, %RMSE=16.35, P<0.0001, Fig. 4G), cellulose (R^2^=0.42, components=4, %RMSE=19.3, P<0.0001, Fig. 4H), lignin (R^2^=0.35, components=3, %RMSE=20.75, P<0.0001, Fig. 4I) and cell solubles (R^2^=0.35, components=4, %RMSEP=19.56, P<0.0001), as well as LMA (R^2^=0.44, N=191, components=3, %RMSE=19.05, P<0.0001) show that that traits could be reliably mapped in Wood River using AISA data (Table S2).

### Detecting plant diversity from remotely sensed spectra

We calculated spectral diversity from airborne data as the mean of the vector normalized spectral distances among all pixels per plot (or subplot), a metric that does not require site-specific model development. At Wood River, based on AISA Kestrel data at the plot scale, spectral diversity readily predicted all measures of plant diversity, including species richness (R^2^=0.497, P< 0.0001, Fig. 4J), phylogenetic species richness (R^2^=0.342, P<0.003, Fig. 4K), and leaf level functional diversity [FD(qDTM)] (R^2^=0.358, P<0.002, Fig. 4L). These relationships did not hold at the subplot scale, however, perhaps because the number of pixels per subplot (6) was insufficient to capture the variability or because there was limited variability at this scale, or both. In BioDIV, spectral diversity based on 0.9 m spatial resolution AVIRIS NG data did not predict any metric of plant diversity, including measured species richness. Prior studies at this site have shown that accuracy in predicting plant diversity is highly dependent on spatial resolution and the high soil fraction/vegetation cover fraction poses difficulties for predicting plant diversity from remotely sensed measures of spectral diversity at a spatial resolution of ∼1 m (Wang et al. 2018, Gholizadeh et al. 2018; Gamon et al. 2020).

Remotely sensing the biodiversity-productivity relationship in either experiment was challenging due to the inability to predict plant diversity metrics from airborne data in BioDIV, and due to the relatively weak measured associations between plant diversity and biomass in Wood River at either the subplot (Fig. 3 D-F) or plot scale (Fig. 3 G-I). Nevertheless, there was a weak relationship between remotely sensed spectral diversity and remotely sensed biodiversity at the subplot scale (Fig. S3C), although this was not significant at the plot scale (Fig. S3B).

Remotely sensed spectral diversity at the plot scale was strongly associated with foliar nitrogen concentration, which may indicate that in ecosystems that are more diverse, plants are able to acquire more nutrients from the soil and allocate it to leaves (Williams et al. 2020), but this same pattern did not emerge at the subplot scale (Fig. S3 D-E).

### Associations between total aboveground inputs and belowground processes and attributes

In the BioDIV experiment, the quantity of total aboveground inputs—or productivity, measured as annual aboveground biomass or vegetation cover—but not their quality, measured as the chemical or phylogenetic-functional composition, were strongly and positively associated with hydrolytic enzyme activity but not oxidative enzyme activity. Vegetation cover and biomass were also positively associated with microbial biomass carbon and nitrogen, net nitrogen mineralization rate, cumulative respiration rate and soil carbon concentration, but not soil nitrogen concentration (Fig. 5; Table S3). Consequently, remotely sensed vegetation cover significantly predicted cumulative soil respiration rates, hydrolytic enzyme activity, microbial biomass carbon, net nitrogen mineralization rates and soil carbon concentration (Fig. 6A-E). In the Wood River experiment, the quantity of total aboveground inputs—measured directly as aboveground biomass or remotely sensed based on PLSR models—were not associated with any of the measured belowground processes and attributes (Fig. 5). In contrast, foliar nitrogen concentration measured on the ground (Table S3) or mapped from remotely sensed data significantly predicted microbial biomass, hydrolytic enzyme activity, net nitrogen mineralization rate, and total soil nitrogen and carbon (Fig. 6). Foliar nitrogen concentration was not associated any of these soil characteristics in the BioDIV experiment (Fig. 5, Table S3).

**Figure 5.**
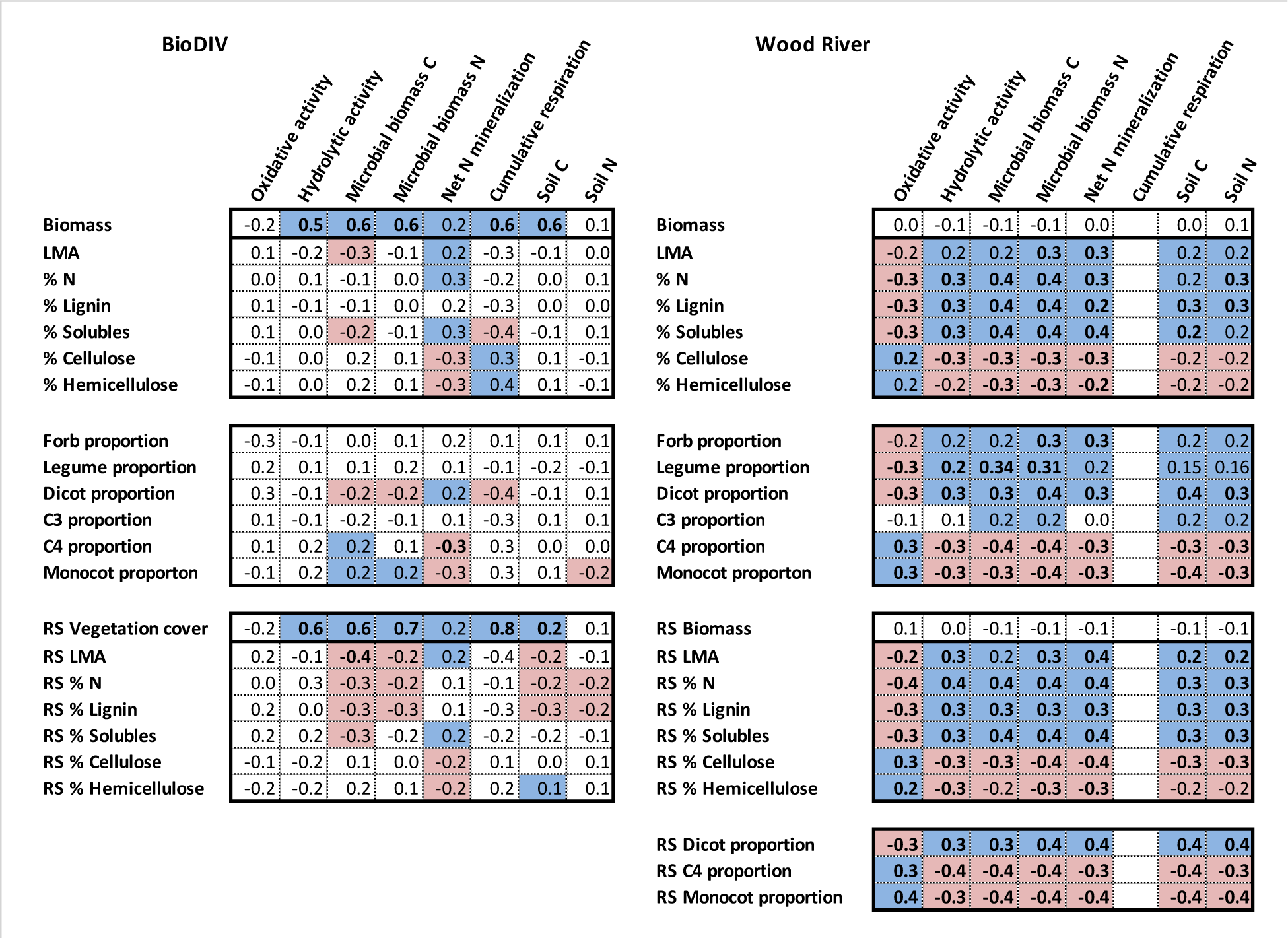
Belowground processes tended to be predicted by remotely sensed measures of biomass in BioDIV but by remotely sensed measures of foliar function and chemical composition in Wood River. Shown are correlation coefficients (r values) at the plot- (BioDIV) or subplot-level (Wood River) between remotely sensed (RS) traits, directly measured leaf traits calculated as community (plot) weighted means, or functional group composition (proportion or biomass) and soil processes, including oxidative enzyme activity (nmol g^-1^ h^-1^), hydrolytic enzyme activity (nmol g^-1^ h^-1^), microbial biomass carbon (mg C [g soil]^-1^), nitrogen net mineralization rate (mg N [g soil]^-1^ d^-1^), cumulative respiration rate (mg CO2-C [g soil^-1^ d^-1^]), and soil carbon and nitrogen concentrations (%). Shading indicates the correlation is significant at P < 0.05 or if bolded at P <0.001. Blue shading indicates the correlation is positive, pale red indicates the correlation is negative. No shading indicates that we found no relationship between the variables, and blank squares indicate no data.

**Figure 6.**
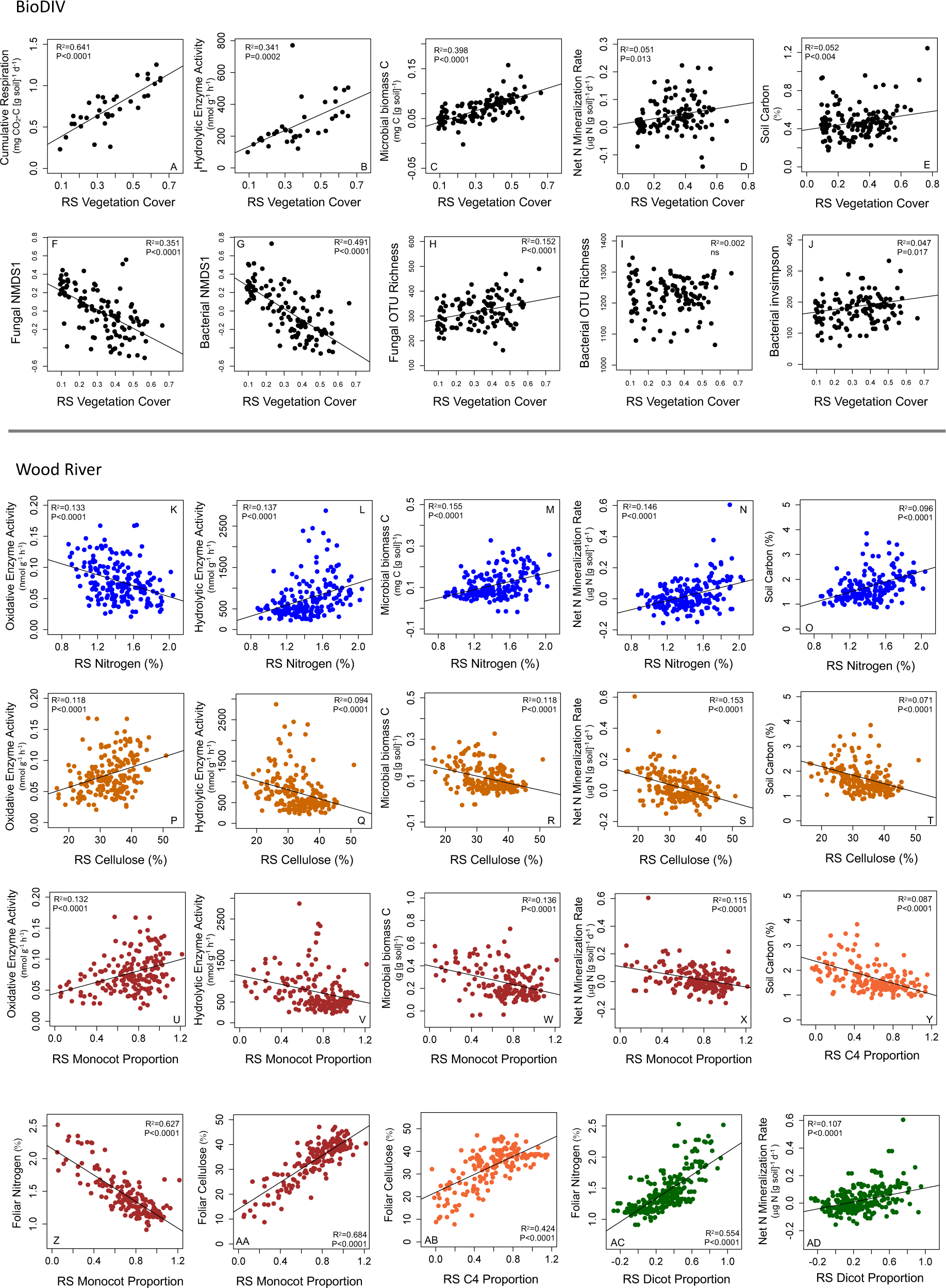
In the BioDIV experiment, remotely sensed vegetation cover predicted A) cumulative respiration rate, B) hydrolytic enzyme activity, C) microbial biomass carbon, D) nitrogen net mineralization rate, E) soil carbon, F) the first NMDS1 axis of the fungal community data and G) the first NMDS1 axis of the bacterial community data as well as H) fungal operational taxonomic unit (OTU) richness but not I) bacterial OTU richness. J) Bacterial diversity measured by the inverse Simpson metric is weakly predicted. Data are from 2015, N=119. In the Wood River experiment, remotely sensed vegetation nitrogen concentration in the vegetation (RS Nitrogen) predicted K) oxidative enzyme activity, L) hydrolytic enzyme activity, M) microbial biomass carbon, N) net nitrogen mineralization rate and O) soil carbon. Remotely sensed (RS) cellulose concentration and RS monocot proportion positively predicted P,U) oxidative enzyme activity, and negatively predicted Q,V) hydrolytic enzyme activity, R,W) microbial biomass carbon, S,X) net nitrogen mineralization rate and T,Y) soil carbon. RS monocot proportion was negatively associated with (Z) foliar nitrogen (%) and positively associated with foliar cellulose concentration (%), as was C4 grass proportion, while dicot proportion was positively associated with foliar nitrogen and soil net nitrogen mineralization rate.

Plant productivity was strongly associated with soil fungal and bacterial composition and to a lesser degree fungal and bacterial diversity in BioDIV. Remotely sensed vegetation cover significantly predicted the first axis of non-metric multidimensional scaling (NMDS) ordination analyses, for both the fungal (Fig. 6F) and bacterial communities (Fig. 6G). These results indicate that the composition of microbial communities is organized in association with plant productivity (Fig. 6F-J). Remotely sensed vegetation cover weakly but significantly predicted fungal diversity, measured as OTU richness (Fig. 6H), but not inverse Simpson diversity (not shown). Remotely sensed vegetation cover also weakly but significantly predicted bacterial diversity, measured as inverse Simpson diversity (Fig. 6J), but not as OTU richness (Fig. 6I).

### Predicting belowground processes and attributes from leaf chemistry and functional group composition

Compared to the strong and consistent associations between total aboveground inputs and, belowground processes or attributes in the BioDIV experiment, directly measured or remotely sensed functional traits showed relatively few associations (Fig. 5; Table S3).

Directly measured LMA, foliar nitrogen concentration, and foliar concentration of cell solubles—measured at the leaf level—were positively while foliar cellulose and hemicellulose concentration were negatively associated with net nitrogen mineralization. The same was true for remotely sensed LMA and foliar concentration of cell solubles, cellulose and hemicellulose, although remotely sensed nitrogen did not show a significant relationship. Cumulative soil respiration was positively associated with directly measured foliar cellulose and hemicellulose concentration and negatively associated with cell soluble concentration. Microbial biomass carbon was negatively associated with LMA and cell soluble concentration—measured at the leaf level or remotely sensed—and with foliar nitrogen concentration and lignin concentration measured at the leaf level. However, most of the measured belowground processes, including oxidative enzyme activity, hydrolytic enzyme activity, microbial biomass nitrogen and total soil carbon and nitrogen concentration were not associated with any of the functional traits measured at the leaf level. Some remotely sensed traits were negatively associated with soil carbon and nitrogen concentration, including foliar nitrogen and lignin concentration, and remotely sensed foliar hemicellulose concentration was significantly but weakly associated with soil carbon concentration. In general, however, functional traits were not strong or consistent predictors of belowground processes in the BioDIV experiment.

Consistent with these findings, phylogenetic-functional group proportion or biomass explained relatively few belowground processes in BioDIV (Fig. 5; Table S3). Dicot proportion was negatively associated with microbial biomass carbon and nitrogen and cumulative respiration, while monocot and C4 grass proportion were positively associated with microbial biomass carbon and negatively associated with net nitrogen mineralization rate. Biomass of these phylogenetic-functional groups showed trends that were similar to both proportion and total biomass relationships with soil attributes. Monocot and C4 grass biomass positively predicted hydrolytic enzyme activity, microbial biomass carbon and nitrogen, cumulative respiration rate and soil carbon, and negatively predicted net nitrogen mineralization rate, similar to total aboveground biomass (Table S3). Forb biomass strongly predicted microbial biomass nitrogen (but not microbial biomass carbon), and net nitrogen mineralization rate. C3 grass and forb biomass also weakly predicted soil carbon concentration (Table S3).

In contrast to BioDIV, at the Wood River site, foliar and remotely sensed plant functional traits were strongly and consistently associated with all belowground processes and attributes, summarized in Fig. 5 and the strength of the regression coefficients were very similar (Table S3). Specifically, both leaf-level and remotely sensed foliar nitrogen (Fig. 6K-O) were negatively associated with oxidative enzyme activity and positively associated with hydrolytic enzyme activity, microbial biomass carbon and nitrogen, net nitrogen mineralization rate and soil nitrogen and carbon—as were leaf-level and remotely lignin, cell soluble concentration and LMA (Table S3). Both leaf-level and remotely sensed foliar cellulose (Fig. 6P-T) were positively associated with oxidative enzyme activity and negatively associated with microbial biomass carbon and nitrogen, net nitrogen mineralization rate and soil nitrogen and carbon concentrations—as were leaf-level and remotely sensed hemicellulose concentration it (Table S3).

The sign of these relationships was largely mirrored phylogenetic-functional group proportion. Monocot proportion—which is strongly negatively associated with remotely sensed foliar nitrogen (Fig. 6Z) and positively associated with cellulose concentration (Fig. 6AA)—was positively associated with oxidative enzyme activity and negatively associated with hydrolytic enzyme activity, microbial biomass carbon, net nitrogen mineralization rate (Fig. 6P-X), and soil carbon (Table S3). C4 grass proportion showed these same patterns although C3 grass proportion showed weakly contrasting patterns or no relationship at all (Fig. 5). Specifically, C3 grass proportion was positively associated microbial biomass carbon and nitrogen as well as soil carbon and nitrogen concentration, but not any of the other soil processes. Dicot proportion—which is strongly associated with remotely sensed nitrogen concentration (Fig. 6AC) and negatively associated with cellulose concentration—was negatively associated with oxidative enzyme activity, and positively associated with hydrolytic enzyme activity, microbial biomass carbon, net nitrogen mineralization rate (Fig. 6AD). The proportion of variation explained in any of these plant functional trait-belowground process relationships was relatively low, ranging from about 5 to 15 %. However, their consistency is striking in the Wood River system (Fig. 5). The consistency in the relationships between the remote sensing traits and soil variables and the leaf traits and soil variables points to a generality in the overall analyses.

### Structural equation models

Our SEM results confirm the hypotheses illustrated in our conceptual models (Fig. 1) and are consistent with the bivariate relationships in Figs. 5 and 6. The analysis synthesizes these findings and shows that vegetation quantity and quality influenced belowground attributes and processes and their interrelationship in the BioDIV and Wood River experiments differently. In BioDIV, the quantity of aboveground inputs to the soil, measured as aboveground biomass, was an important positive predictor of the activity and abundance of soil microbes, while vegetation quality, measured as the carbon to nitrogen ratio in aboveground vegetation, was not (Fig. 7A). The quantity of aboveground inputs also was positively associated with soil nutrient availability, measured as net nitrogen mineralization rate (Nmin), while vegetation quality was not.

**Figure 7.**
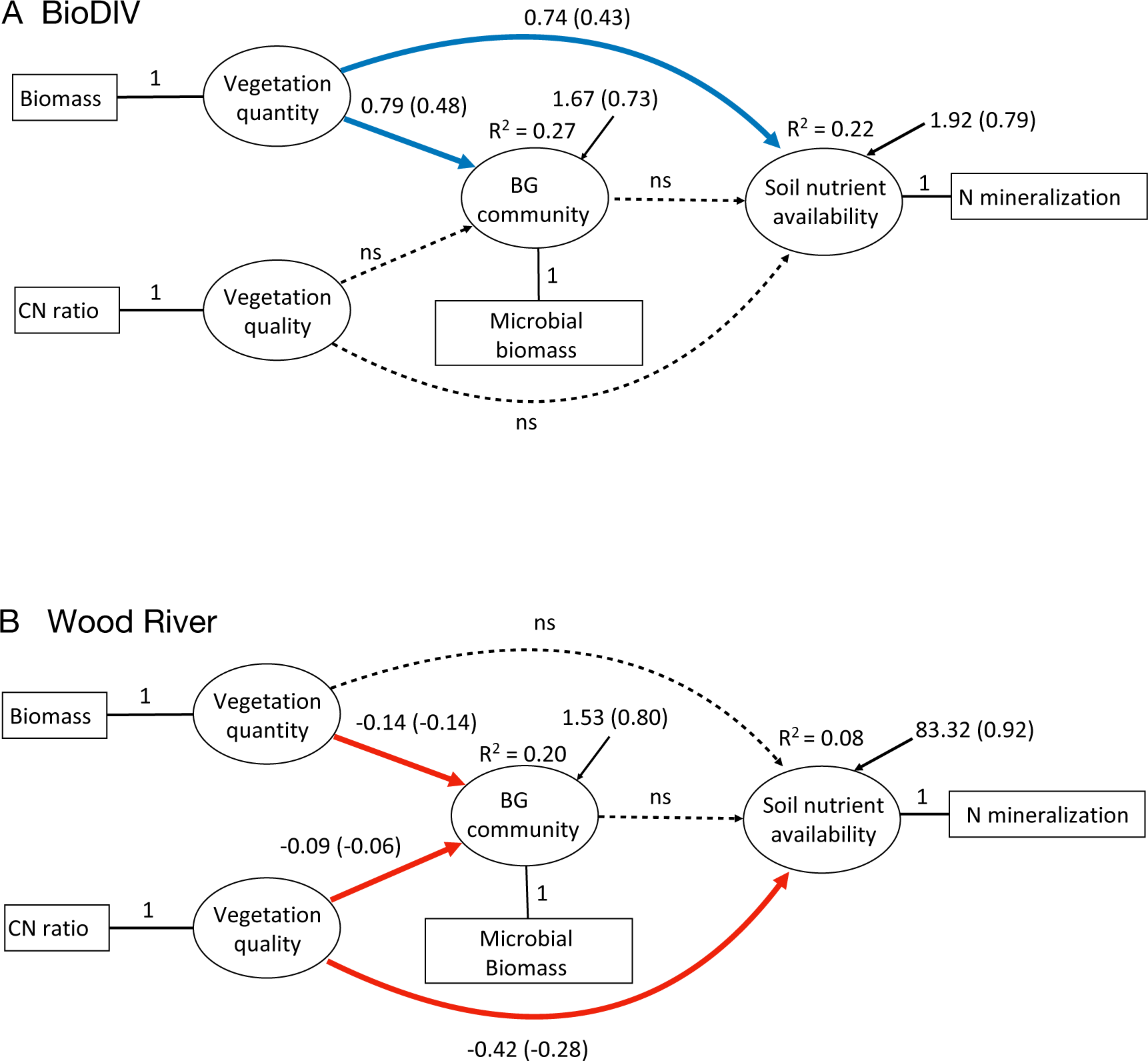
Structural equation models reveal the relative importance of aboveground quantity and quality of vegetation inputs for belowground microbial processes and soil nutrient availability and their interrelations in the BioDIV [Chi-square = 0.30, df = 1, p = 0.59] (A) and Wood River [Chi-square = 0.02, df = 1, p = 0.89] (B) experiments. In each model, the latent variable constructs (ovals) are represented by measured variables (rectangles). Vegetation quantity and quality are represented by aboveground biomass and vegetation carbon to nitrogen ratio, respectively. The latent construct of belowground microbial community functions is represented microbial biomass carbon, while the soil nutrient availability is represented by net nitrogen mineralization rate. Blue arrows indicate significant positive relationships. Red arrows indicate significant negative relationships. Dashed lines non-significant relationships. Coefficients and standardized coefficients (in brackets, one unit change in the predictor variable causes the amount of one standardized coefficient change in the dependent variable) are given next to the arrows. Predictor variables were converted the following way: one unit of biomass equals biomass g m^-2^/100, one unit nitrogen net mineralization (Nmin) rate (mg N [g soil]^-1^ d^-1^) equals Nmin*100, and one unit microbial biomass carbon (microbioC) (mg C [g soil]^-1^) equals microbioC*10.

In contrast, in the Wood River system, vegetation quality, measured as the carbon to nitrogen ratio of aboveground vegetation, was strongly and negatively associated with soil nutrient availability, measured as net nitrogen mineralization rate (Nmin) (Fig. 7B), while the quantity of aboveground inputs to the soil, measured as aboveground biomass, was not. In addition, in this system both vegetation quality and vegetation quantity, measured as aboveground biomass, was negatively associated with the abundance of microbes in the soil, such that greater amounts of vegetation nitrogen relative to carbon increased microbial biomass.

## Discussion

Belowground biodiversity and processes are critical to essential ecosystem functions such as soil fertility and carbon sequestration that maintain the Earth’s life support systems. Yet they are difficult to study across large spatial scales. Here we demonstrate potential for using spectroscopic imagery across broad spatial extents to predict belowground properties and functions in grassland systems that varied in diversity and productivity. In a relatively low productivity system, with low soil carbon and nitrogen concentrations (BioDIV), the supply of substrates and thus total vegetation biomass limited microorganisms and their functions in the soil, influencing microbial community composition and nutrient dynamics. In a more productive system with higher soil carbon and nitrogen concentrations (Wood River), where substrate supply was less limiting to soil microbes, the chemical composition of vegetation had a predominate influence on microbial processes and nutrient dynamics.

The IPBES 2018 Americas Assessment documented that 35% of grassland systems in the Americas have been converted to pasture or grazing lands and only a small fraction of intact grassland systems remains (Cavender-Bares et al. 2018). In North America alone, a large fraction of grasslands has been altered and their ecosystem functions degraded relative to pre-European settlement, reducing their productivity. The range of grassland systems—from those more heavily altered by humans to those that are relatively intact— has been simulated to some extent by these diversity experiments in grassland systems, both at the low end of a diversity-productivity gradient (BioDIV in central Minnesota) and towards the higher end of a diversity-productivity gradient (Wood River in central Nebraska).

In such experiments, we show that remotely sensed variables of prairie grassland ecosystems related to both the quantity and quality of aboveground inputs predicted belowground processes. Airborne spectroscopic data at 0.9 m or 1 m resolution predicted aboveground biomass, foliar chemistry and function, and where vegetation cover was high, phylogenetic-functional group composition. These aboveground metrics, in turn, predicted belowground microbial and soil processes, as well as nutrient availability in both systems. However, the relative importance of the total quantity of vegetation biomass—or total energy inputs—compared to the functional and chemical quality of these inputs in driving microbial communities and belowground processes varied between the two experimental systems. In BioDIV, the quantity of vegetation inputs to the soil was the most critical predictor for microbial abundance and soil processes, while the quality of inputs was of secondary importance. The greater importance of quantity over quality of aboveground inputs likely occurred because of high variation in productivity and the very low productivity associated with low diversity levels in BioDIV. Weeding to maintain the low diversity treatments greatly reduces plant cover causing the low diversity plots to have very low productivity compared to most grasslands—even compared to heavily influenced grasslands (Hui & Jackson, Knapp et al. 2001; Fay et al. 2015). In Wood River, which had a higher plant density resembling more typical prairie, the quality of vegetation inputs was the most important predictor for microbial abundance and soil processes.

In grassland systems, detecting taxonomic composition and diversity levels at pixel sizes of ∼1 m or greater remains complicated due to a mismatch between pixel size and the size of individual plants (Wang et al. 2018, Gholizadeh et al. 2018). While plant biodiversity was readily predicted by spectral diversity from airborne imagery at Wood River (Fig. S2, Gholizadeh et al. 2019), accurate detection in BioDIV was possible only with very high spatial resolution (<10 cm) (Wang et al., 2018) using a ground-based robotic tram (Gamon et al., 2006). Using a different metric of spectral diversity—the mean distance between vector-normalized spectra among pixels—we again find at Wood River that spectral diversity predicts plant diversity, including species richness, phylogenetic species richness (PSR) and leaf-level functional diversity (qDTM) at the plot scale but not at the subplot scale. Remotely sensed spectral diversity was not significantly associated with remotely sensed biomass at the plot scale (Fig. S3B), and directly measured metrics of plant diversity were only weakly associated at this scale (Fig. 3 G-I). At the subplot scale these relationships were significant—both directly measured (Fig. 3 D-F) and remotely sensed (Fig. S3C)—but spectral diversity explained a low proportion of the variance in biomass. In BioDIV, where the relationships between measures of plant diversity and productivity are stronger, remotely sensed spectral diversity did not predict metrics of plant diversity at 1 m resolution. Nevertheless, Schweiger et al (2018) found that spectral diversity, measured both at the leaf level and remotely using 1 cm tram data, accurately predicted plant functional diversity and phylogenetic diversity in BioDIV, and that spectral diversity strongly predicted biomass. These results highlight the need for close attention to spatial scale – both in terms of grain size of measurement and consideration of the spatial extent at which relationships between biodiversity and ecosystem functions can be observed (Gamon et al. 2020).

The ability to remotely map functional traits and recover the same relationships with soil processes as measured trait values demonstrates promise for current and forthcoming satellite missions and airborne detection efforts of a range of ecosystem processes through mapping of functional traits with spectroscopy. Where canopy conditions permit—which we hypothesize to depend on high vegetation cover— mapping of phylogenetic-functional groups is also likely to be predictive of belowground processes. Applying this approach to other systems will require an understanding of the extent to which substrate quantity limits microbial processes and the relative importance of vegetation quality in driving belowground dynamics. Vegetation cover and biomass can be remotely sensed with high accuracy using imaging spectroscopy and have the potential to inform where on the diversity-productivity continuum an ecosystem resides, and thus whether total substrates are a limiting factor or whether chemical and phylogenetic-functional groups are likely to be more predictive of belowground processes and attributes. Targeted field campaigns to test the expected relationships between vegetation inputs and belowground processes, informed by cover, biomass and trait maps, will be important.

### Relationships between vegetation cover, productivity and belowground attributes and processes

Total organic matter inputs to soils are likely much lower in BioDIV than in Wood River, for several reasons, and thus likely to limit soil microbes at BioDIV. In BioDIV, the diversity treatments (1-16 species/plot, drawn from a pool of 32) create 16-fold variation in biomass (25 - 283 g m^-2^ in 2015) and hence in the quantity of substrate inputs to the soil. Yet even the most diverse and productive plots have modest productivity relative to the original tall grass prairie of the region. At Wood River, with higher species richness/plot and a total species pool of ca. 80 species, biomass ranges from 285 - 1079 g m^-2^ (in 2017), such that the least productive communities at this site are comparable to the most productive communities in BioDIV. While root turnover and rhizodeposition provide an important source of substrate inputs to the soil in both systems, aboveground litter inputs are likely a greater contribution to total organic matter inputs to soils at Wood River, given more frequent burning of aboveground biomass in BioDIV than at Wood River (annually or biannually vs. once every five years). In addition, higher productivity at Wood River likely arises in part from a warmer climate, finer-textured soils with deposits from the Platte River, and thus higher soil fertility.

Lower productivity at BioDIV for reasons mentioned above likely lead to stronger microbial substrate limitation there than at Wood River. Previous studies have documented that when substrate quantity is limiting, it is positively related to microbial biomass and fungal abundance (Wardle, 1992; Zak et al., 1994; Cleveland & Liptzin, 2007; Whitaker et al., 2014) and to soil carbon, which has been previously documented in BioDIV (Fornara & Tilman, 2008, Yang et al. 2019). Higher microbial biomass and soil carbon are thus expected to be associated with high soil respiration (Whitaker et al., 2014). In systems where energy inputs primarily limit microbial processes, nutrient composition of soil inputs is secondary (Hättenschwiler & Jørgensen, 2010; Whitaker et al., 2014; Cline et al., 2017). A similar pattern appears to be the case in BioDIV, where chemical composition of the above ground vegetation is relatively less important and less predictive than total energy inputs to microbial abundance, composition, activity and the soil processes they drive (Zak et al., 2003).

### Relationships between vegetation chemistry, foliar functional traits, phylogenetic-functional groups and belowground attributes and processes

Phylogenetic-functional group composition—associated with changes in vegetation chemistry and root chemistry—was a stronger influence on microbial composition, biomass and activity in Wood River than in BioDIV. In the Wood River system, remotely sensed foliar chemistry and plant functional-phylogentic group composition, as well as in situ measures of these same traits and phylogenetic-functional group proportion, were stronger predictors of belowground enzyme activity, net nitrogen mineralization rates, microbial biomass, and soil carbon and nitrogen compared to BioDIV. Nevertheless, the proportion of the major phylogenetic lineages—monocots and dicots—as well as individual functional groups, particularly C4 grasses, to varying extents remained predictors of microbial biomass carbon and nitrogen, cumulative soil respiration and net nitrogen mineralization rates in BioDIV, as did some aspects of chemical composition, consistent with past studies at BioDIV and the nearby BioCON experiment (Dijkstra et al. 2006; Fornara et al., 2009; Mueller et al. 2013; Wei et al. 2019).

In Wood River, oxidative enzyme activity was negatively predicted by leaf level and remotely sensed LMA, foliar nitrogen and lignin concentration, and foliar cell soluble concentration and positively predicted by leaf level and remotely sensed foliar cellulose and hemicellulose concentration. In contrast, hydrolytic enzyme activity, microbial biomass carbon and nitrogen, and soil carbon were positively predicted by foliar nitrogen, cell solubles, lignin and LMA. Foliar nitrogen and lignin concentration are highly correlated in both of these grassland systems, such that teasing apart their independent relationships with other factors is problematic. Nitrogen, both when applied externally and in leaf litter, has been shown to inhibit oxidative enzyme activity (Fenn et al., 1981; Carreiro et al., 2000; DeForest et al., 2004; Zak et al., 2008; Edwards et al., 2011; Hobbie et al., 2012), which may explain why these enzymes show reduced activity in plots with higher foliar nitrogen despite higher lignin inputs. Greater soil carbon is expected to support higher microbial biomass (Wardle, 1992; Zak et al., 1994; Cleveland & Liptzin, 2007), which helps explain higher hydrolytic enzyme activity in plots with higher foliar nitrogen (Whitaker et al., 2014).

Nitrogen in substrates can control the composition and activity of microbial communities and the processes they drive (Hättenschwiler & Jørgensen, 2010). If litter- feeding organisms and decomposer communities are limited by the relative availability of nitrogen, their activity is expected to increase with substrates higher in N. We found that both remotely sensed foliar nitrogen concentration as well as the proportion of legumes positively predicted microbial biomass carbon and nitrogen and their hydrolytic enzyme activity and nitrogen net mineralization rates, consistent with results from BioCON, adjacent to BioDIV (Dijkstra et al. 2006; Mueller et al. 2013; Wei et al. 2019). Likewise, the proportion of C4 grasses and of grasses, in general—which have low tissue N concentrations and high cellulose and hemicellulose concentrations—was a negative predictor of these same processes. These results demonstrate that remotely sensed traits mapped at landscape scales as well as phylogenetic-functional group composition can be predictive of belowground processes and attributes. However, in situ characterization of the systems is necessary to understand how vegetation inputs drive soil processes—as well as to determine the extent to which traits or phylogenetic-functional groups can be robustly mapped.

### Relevance of phylogenetic-functional groups for predicting belowground processes and attributes using imaging spectroscopy

Our results show that airborne data can provide critical information relevant to belowground ecosystem processes, both in terms of quantity and quality of inputs to soil, that are not easily obtained from the ground. Chemical sampling of vegetation is time- consuming and costly, particularly over large spatial extents. Airborne information, coupled with increasingly available vegetation trait and biomass models, will eventually provide this kind of information with ultimately lower cost and less time. However, our work shows that modeling how the quality and quantity of inputs influence belowground processes requires characterizing systems in advance. The relative importance of total aboveground inputs to soil processes compared to the chemical composition of those inputs is expected to vary along the diversity-productivity gradient, given that carbon is not likely to be limiting at the high productivity end of this gradient. Understanding where a given ecosystem falls along the plant diversity-productivity gradient is likely critical for determining whether remotely sensed measures of productivity or vegetation chemistry are most predictive of belowground microbial processes and soil nutrients.

Based on our results from two contrasting experimental grassland systems, we can anticipate that belowground processes of overgrazed and degraded grasslands that are low in biodiversity and productivity may be driven by input quantity. In contrast, belowground processes in less disturbed prairie systems may be more driven by vegetation composition and quality. If these relationships can be established generally at large spatial scales across grassland systems, we will be in a position to advance understanding of belowground ecosystem processes in these globally threatened ecosystems, facing land use change, increasing grazing pressure and a range of other anthropogenic factors.

## Acknowledgements

The project was funded by the Dimensions of Biodiversity program through NSF/NASA DEB-1342872; NSF DBI: 2021898 and the Cedar Creek Long Term Ecological Research Program through NSF DEB 1831944. We thank Ian Carriere, Brett Fredericksen, Jesús Pinto-Ledezma, Chris Buyarski, Troy Mielke, Shan Kothari, and numerous Cedar Creek research interns for field and laboratory assistance, Laura Williams for assistance with scripting and Lauren Cline for organization of and access to fungal and soil data in BioDIV. We note that a revised version of this manuscript has been accepted by *Ecological Monographs* but is not yet available for distribution.

