## Supplementary material for "Remotely detected plant function in two midwestern prairie grassland experiments reveals belowground processes": SI1.Supplemental Figures

**Appendix S1**

Cavender-Bares, J. M., A. K. Schweiger, J. A. Gamon, H. Gholizadeh, K. Helzer, C. Lapadat, M. D. Madritch, P. A. Townsend, Z. Wang, and S. E. Hobbie. Remotely detected plant function in two midwestern prairie grassland experiments reveals belowground processes

**Figure S1.**
Two grassland biodiversity experiments. Left: The BioDIV experiment at Cedar Creek in Minnesota, photo: Jacob Miller 2014. Right: The Nature Conservancy Wood River diversity experiment in Nebraska, photo: Chris Helzer 2018. In the BioDIV experiment, 150 maintained plots are 81 $m^2$, and the number of species planted per plot varies from 1 - 16 with a range of total biomass from 10 - 60 g $m^{-2}$. In the Wood River experiment, 24 large plots, each approximately 3600 $m^2$, were planted with 11 - 48 species and sampled in 6 $m^2$ subplots along two transects (192 total subplots). Biomass ranges from 280 to 1100 g $m^{-2}$. Inset: Stars on map show approximate locations of the two experiments with BioDIV northeast of Wood River in the midwestern U.S.

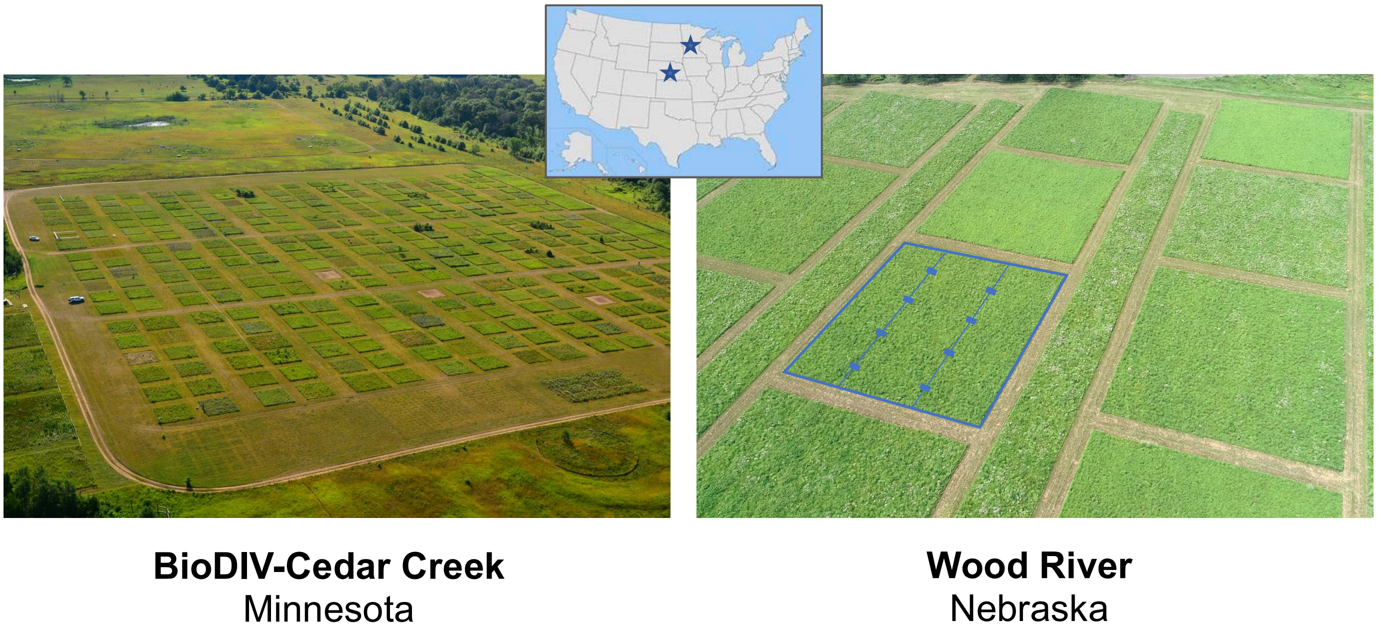

**Figure S2**
Correlation matrix for chemical and functional traits measured at the leaf level on species in the Wood River experiment and biomass per unit area in the experimental plots. Traits include concentrations of hemicellulos, cellulose, lignin, nitrogen, leaf mass per area, cell solubles and carbon. Blue ellipses indicate positive associations and red ellipses show negative associations. Narrower ovals and darker colors indicate stronger relationships, as indicated by the r value scale to the right of the graph.

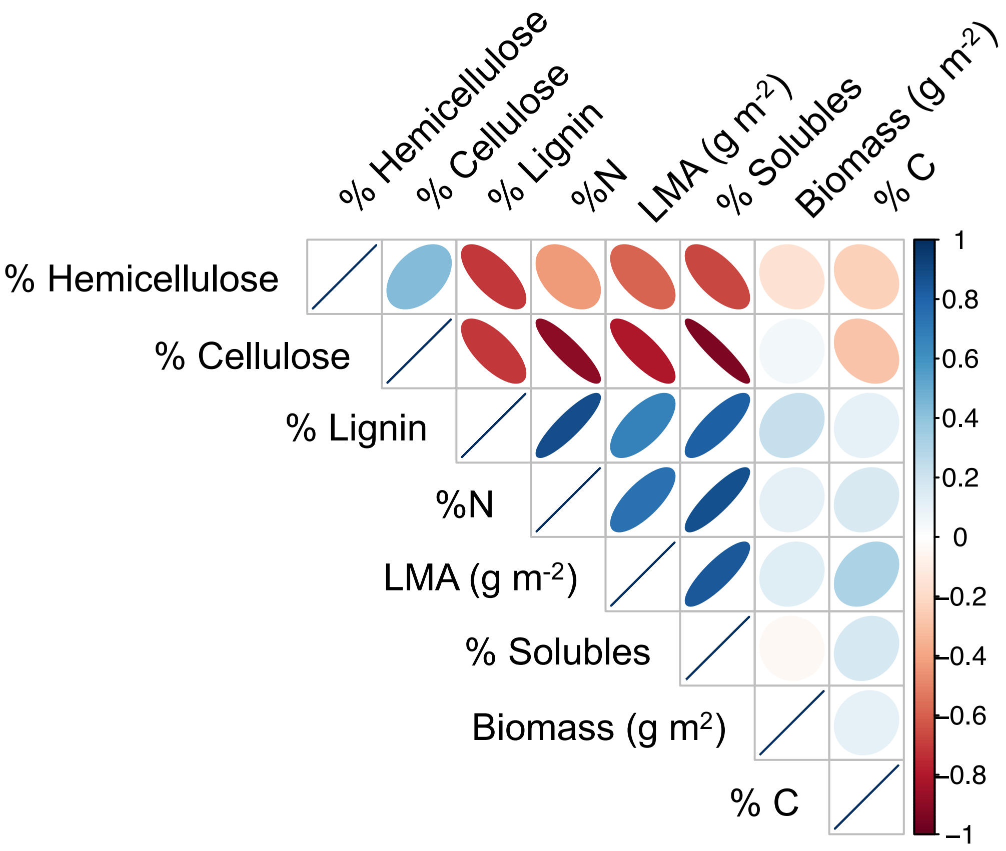

**Figure S3**

Aboveground plant biomass predicts belowground root biomass, BioDIV 2015, with belowground biomass often five-fold greater than aboveground biomass (A). In BioDIV, vegetation cover (%) consistently predicts aboveground plant biomass in 2014 (B), 2015 (C) and 2016 (D). In contrast, the Wood River experiment has 100% vegetation cover in all of the plots and vegetation cover does not predict aboveground plant biomass (E).

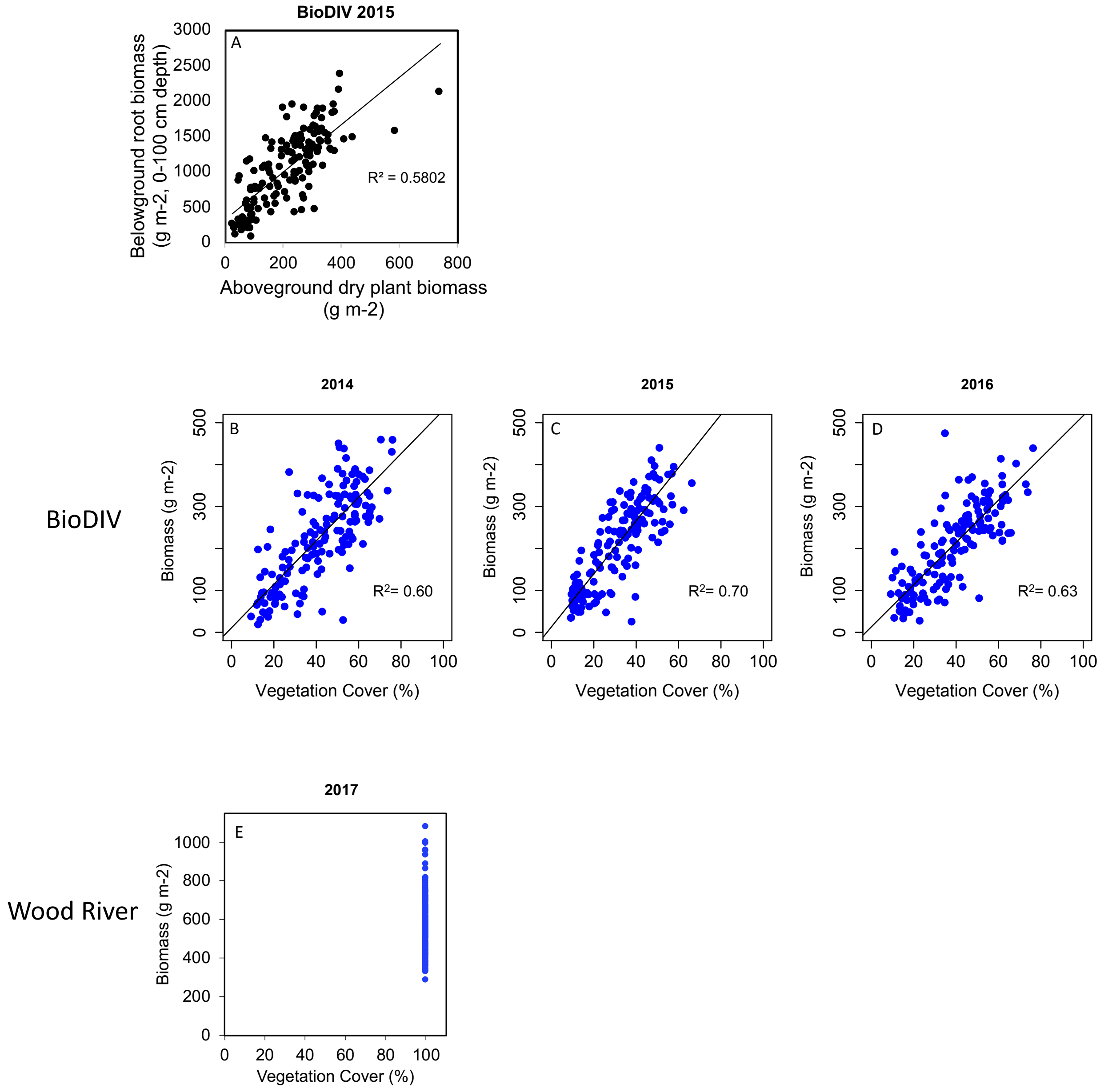

**Figure S4**

Relationships between A) soil carbon concentration (%) and plant biomass (g m$^{-2}$), B) microbial biomass carbon (mg C [g soil]$^{-1}$) and soil carbon concentration, and C) microbial biomass carbon and plant biomass in the BioDIV experimental plots (black circles) and the Wood River experimental subplots.

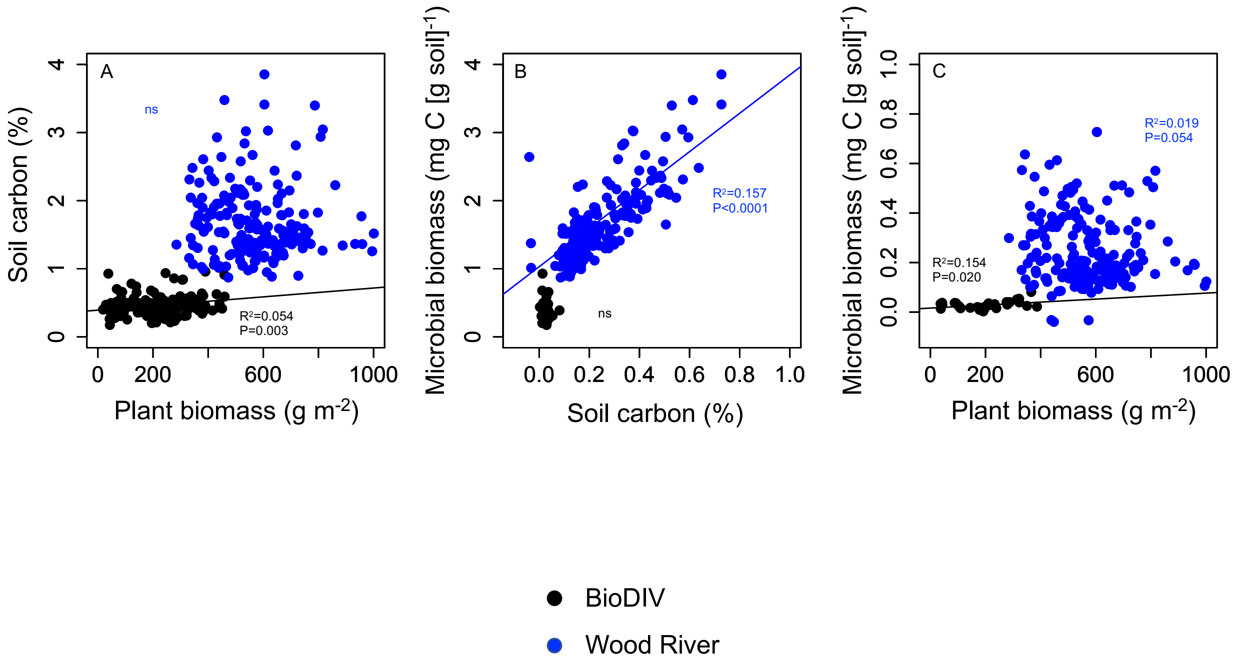

**Figure S5**

Relationships between remotely sensed biomass, spectral diversity and nitrogen concentration at Wood River. A) Relationship between the predicted biomass—averaged for all pixels at the plot scale based on PLSR models trained from data at the subplot scale—and the measured biomass averaged per plot. B) Remotely sensed predicted biomass for all plots in relation to remotely sensed spectral diversity, calculated as the mean vector normalized spectral distances among all pixels in the plot. C) Remotely sensed predicted biomass as the subplot scale in relation to remotely sensed spectral diversity. D) Remotely sensed vegetation nitrogen concentration in relation to spectral diversity at the plot scale and the E) subplot scale.

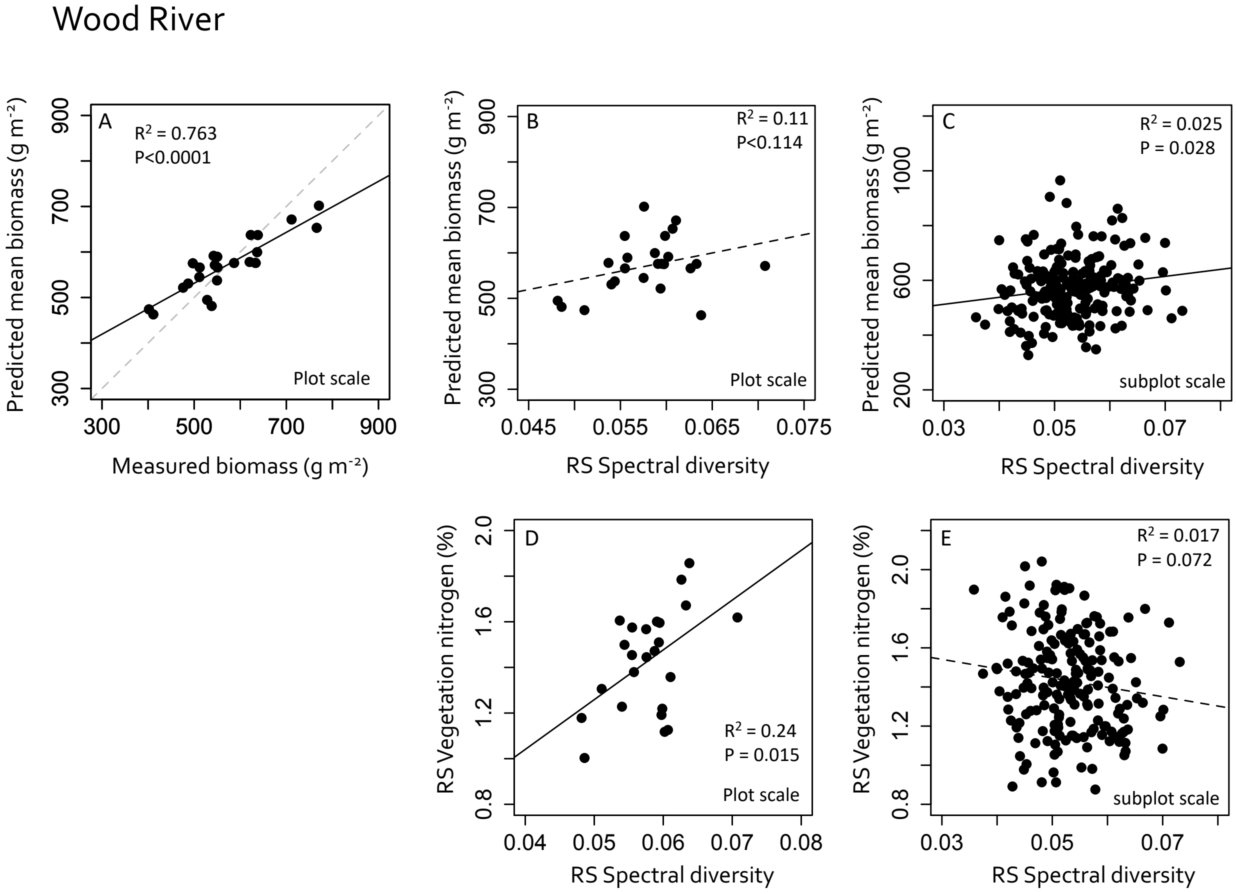
