## Supplementary material for "Remotely detected plant function in two midwestern prairie grassland experiments reveals belowground processes": SI2.Supplemental Tables

**Appendix S2**

Cavender-Bares, J. M., A. K. Schweiger, J. A. Gamon, H. Gholizadeh, K. Helzer, C. Lapadat, M. D. Madritch, P. A. Townsend, Z. Wang, and S. E. Hobbie. Remotely detected plant function in two midwestern prairie grassland experiments reveals belowground processes

**Table S1**. List of data collected in the BioDIV experiment at the Cedar Creek Ecosystem Science Reserve in Minnesota and in the Wood River Nature Conservancy prairie diversity experiment in Nebraska, including year collected, and whether the data component has been previously been published.

| Year | Experiment | Factor measured on the ground | Remotely sensed data collected | Number of plots (or subplots) | Previously published |
|---|---|---|---|---|---|
| 2014 | BioDIV | | AVIRIS NG | All plots | Wang et al 2019 |
| 2014 | BioDIV | | Plant trait map | All plots | Wang et al 2019 |
| 2014 | BioDIV | | Plant FD | All plots | Current study |
| 2014 | BioDIV | Plant aboveground biomass | | 154 plots | Cline et al 2018 |
| 2014 | BioDIV | Plant richness and species biomass | | 154 plots | Cline et al 2018 |
| 2014 | BioDIV | Functional group biomass and proportion | | 154 plots | Cline et al 2018 |
| 2014 | BioDIV | Functional diversity - leaf level | | | Schweiger et al 2018 |
| 2014 | BioDIV | Soil extracellular enzyme activity | | 35 plots | Cline et al 2018 |
| 2014 | BioDIV | Microbial biomass | | 35 plots | Cline et al 2018 |
| 2014 | BioDIV | Soil N and C | | 154 plots | Cline et al 2018 |
| 2015 | BioDIV | Leaf level spectral data | | | Schweiger et al 2018 |
| 2015 | BioDIV | Functional diversity - leaf level | | | Schweiger et al 2018 |
| 2015 | BioDIV | Bacterial composition | | 35 plots | Current study |
| 2015 | BioDIV | Bacterial diversity | | 35 plots | Current study |
| 2015 | BioDIV | Fungal composition | | 35 plots | Cline et al 2018 |
| 2015 | BioDIV | Fungal diversity | | 35 plots | Cline et al 2018 |

| | | | | | |
|---|---|---|---|---|---|
| 2015 | BioDIV | | AVRIS NG | all plots | Wang et al 2019 |
| 2015 | BioDIV | Root chemistry | | 35 plots | Current study |
| 2015 | BioDIV | Soil respiration | | 35 plots | Cline et al 2018 |
| 2015 | BioDIV | Net mineralization rate | | 35 plots | Cline et al 2018 |
| 2016 | BioDIV | | AVIRIS NG | all plots | Wang et al 2019 |
| 2016 | BioDIV | Soil N and C | | 154 plots | Current study |
| 2017 | Wood River | | CALMIT-AISA Kestrel | all plots | Gholizadeh et al 2019 |
| 2017 | Wood River | | Plant trait maps | | Current study |
| 2017 | Wood River | Plant richness in plots | | 36 plots | Gholizadeh et al 2019 |
| 2017 | Wood River | Plant species percent cover subplots | | 144 subplots | Current study |
| 2017 | Wood River | Plant aboveground biomass | | 144 subplots | Current study |
| 2017 | Wood River | Leaf level spectral data | | 144 subplots | Current study |
| 2017 | Wood River | Leaf level functional diversity | | 144 subplots | Current study |
| 2017 | Wood River | Soil extracellular enzyme activity | | 144 subplots | Current study |
| 2017 | Wood River | Microbial biomass | | 144 subplots | Current study |
| 2017 | Wood River | Net nitrogen mineralization rate | | 144 subplots | Current study |
| 2017 | Wood River | Soil NO3, NH4, N and C | | 144 subplots | Current study |
