## Supplementary material for "Remotely detected plant function in two midwestern prairie grassland experiments reveals belowground processes": SI3.Supplemental Methods

**Appendix S3**

Cavender-Bares, J. M., A. K. Schweiger, J. A. Gamon, H. Gholizadeh, K. Helzer, C. Lapadat, M. D. Madritch, P. A. Townsend, Z. Wang, and S. E. Hobbie. Remotely detected plant function in two midwestern prairie grassland experiments reveals belowground processes

### Bacterial DNA analysis methods

Bacterial DNA as extracted using the Fas-tDNA SPIN Kit (MP Biomedical, Solon, Ohio, USA). The 16S rRNA genes were amplified using primers 515f and 806r (Caporaso et al. 2011). PCR reactions contained 1 µL of ~20 ng/µL DNA template, 1 µL of each 5 µM primer, 12 µL of Nuclease-Free water (Quiagen, Hilden, Germany), and 10 µL Q5 High-Fidelity 2X Master Mix (New England BioLabs, Ipswich, Massachusetts, USA) to yield 25 µL reaction mixtures. Bacterial PCR cycling parameters were 94°C for 180 s, followed by 35 cycles of 94°C for 45 s, 55°C for 60 s, and 72°C for 90 s, followed by 600 s extension at 72°C. Positive and negative controls were included with each PCR reaction set. All PCR reactions were run in duplicate then combined before quantification with PicoGreen (Invitrogen, Paisley, UK) and again verified on a 1% agarose gel using the Molecular Imager Gel Doc XR system (Bio-Rad Laboratories, Hercules, California, USA). PCR products were combined in equimolar concentrations and the composite sample was gel-purified on a 1% agarose gel before sequencing at the West Virginia Genomics Core Facility in Morgantown, West Virginia, for sequence analysis with the Illumina MiSeq platform (Illumina, San Diego, California, USA) with 250 paired-end reads at West Virginia University's Genomic Core Facility. Bacterial Illumina MiSeq pair-end reads were processed using MOTHUR version 1.35.1 using the MOTHUR MiSeq standard operating procedure (Schloss et al. 2011; Kozich et al. 2013). We screened joined sequences and removed any sequence that was longer than 275bp, contained at least one ambiguous base, and/or had >8 nt homopolymers. Sequences were further screened for singletons. Aligned and screened sequences were clustered at 97% similarity and operational taxonomic unit (OTU) classification of unique sequences was completed using the Ribosomal Database Project (RDP) taxonomic database (release 9; Cole et al. 2009). To account for variation in the number of sequence reads per sample, the number of sequence reads in each sample was rarefied to 52,880, which removed one sample from further bacterial analysis due to poor sampling depth.
